# Plant-associated fungi co-opt ancient antimicrobials for host manipulation

**DOI:** 10.1101/2024.01.04.574150

**Authors:** Fantin Mesny, Valentina Wolf, Ana López-Moral, Anton Kraege, Wilko Punt, Jiyeun Park, Jinyi Zhu, Yukiyo Sato, Bart PHJ Thomma

## Abstract

Evolutionary histories of effector proteins secreted by fungal pathogens to mediate plant colonization remain largely elusive. While most functionally characterized effectors modulate plant immunity, recent discoveries have revealed novel functions in targeting host-associated microbiota. We now developed an Antimicrobial Activity Predictor for Effector Candidates (AMAPEC), and identified a wealth of antimicrobial effectors, including many highly conserved ones — suggesting ancient evolutionary origins. Surprisingly, several plant immune-modulating effectors display antimicrobial activity. We propose that these evolved from ancestral antimicrobials while retaining their original functions. In addition to roles in suppressing host immunity, they may manipulate plant microbiota to promote colonization. We argue that microbial antagonism is a fundamental fungal effector function and suggest that fungi repurposed ancient antimicrobials to serve multiple roles during host-pathogen co-evolution.

## Introduction

Under continuous threat of microbial parasitism, organisms evolved immune systems to withstand pathogenic encounters (*1–4*). Conversely, pathogens evolved strategies to interfere with immune responses and support host colonization. Plant-pathogenic fungi secrete effectors that target a wide variety of molecular mechanisms *in planta*, resulting in compromised host immunity and resilience (*5, 6*). While well-documented effector targets include plant metabolism and immunity, effectors may also serve in microbial competition, as revealed by the recent identification of effectors with selective antimicrobial properties that suppress antagonistic plant microbiota members with disease-suppressive functions (*7–14*). It is generally accepted that effectors are the products of long co-evolutionary “arms races” in which plants and pathogens aim to defend and overcome defenses, respectively (*1*, *3*). Yet, the evolutionary histories and, more particularly, the molecular origins of most effectors remain enigmatic, although some may originate from gene divergent evolution, genome recombination or horizontal gene transfer during host adaptation (*15–18*).

Here, we investigate evolutionary histories of effectors in the light of the recent discovery of antimicrobial effectors. As we reveal that many secreted antimicrobial proteins show broad conservation throughout the fungal kingdom, suggesting ancient origins, we hypothesize that effectors with functions in host physiology manipulation evolved from ancient antimicrobials. We validate this hypothesis by demonstrating that effectors with reported functions in host immunomodulation have ancestral antimicrobial properties that may serve in microbial antagonism during plant colonization. Thus, our study provides unprecedented insights into fungal evolution and the origins of key factors required for host colonization.

## Results

### Accurate prediction of effector antimicrobial activity

We first sought to identify novel fungal effectors with antimicrobial activities. Machine learning classifiers have previously supported the identification of antimicrobial peptides (AMPs) that are shorter than 100 amino acids in length, based on their physicochemical properties (*19–21*). However, these are not suited to predict antimicrobial activities of effectors, as they are generally longer and form more complex structures (fig. S1). To train an adequate predictor, we curated a dataset of experimentally validated antimicrobial proteins from diverse organisms in the size range of effectors (35–642 amino acids, median = 125.5; Fig. 1, A and B and table S1). Additionally, we curated a negative training set of equivalent proteins unlikely to have antimicrobial activity according to their functional annotation (table S2). Carbohydrate-active enzymes (CAZymes) were not included since, although several families target microbial structures, it is often unknown whether individual members compromise microbial growth and can thus be classified as genuine antimicrobials. For each protein in the training sets, we calculated properties from their amino acid sequences and predicted high-confidence protein structures (Fig. 1, C and fig. S2), representing 70 numerical values reflecting diverse physicochemical properties (table S3). Moreover, we queried for the presence/absence of six k-mers that are over- or underrepresented in the sequences of the antimicrobial proteins (Fig. 1, C and table S3). All data were used to train a Support Vector Machines classifier and subsequently estimated its quality through leave-one-out cross-validation, revealing that our classifier has high accuracy, recall and specificity, particularly for fungal proteins (Fig. 1, D). Analysis of support vector coefficients, representing the importance of individual properties for the prediction, revealed a role for hydrophobicity, charge, secondary structures, identity of exposed amino acids, disulfide bonds and structural cavities (fig. S3). To confirm that our predictor can reliably identify novel fungal antimicrobial proteins, we tested whether it correctly calls out seven more recently characterized fungal antimicrobials that were not included in the training set (*22–26*). As all seven proteins were correctly classified as antimicrobials (table S4), we implemented the predictor in the novel software package AMAPEC (**a**nti**m**icrobial **a**ctivity **p**rediction for fungal **e**ffector **c**andidates; Fig. 1, E), available at https://github.com/fantin-mesny/amapec.

**Fig. 1.**
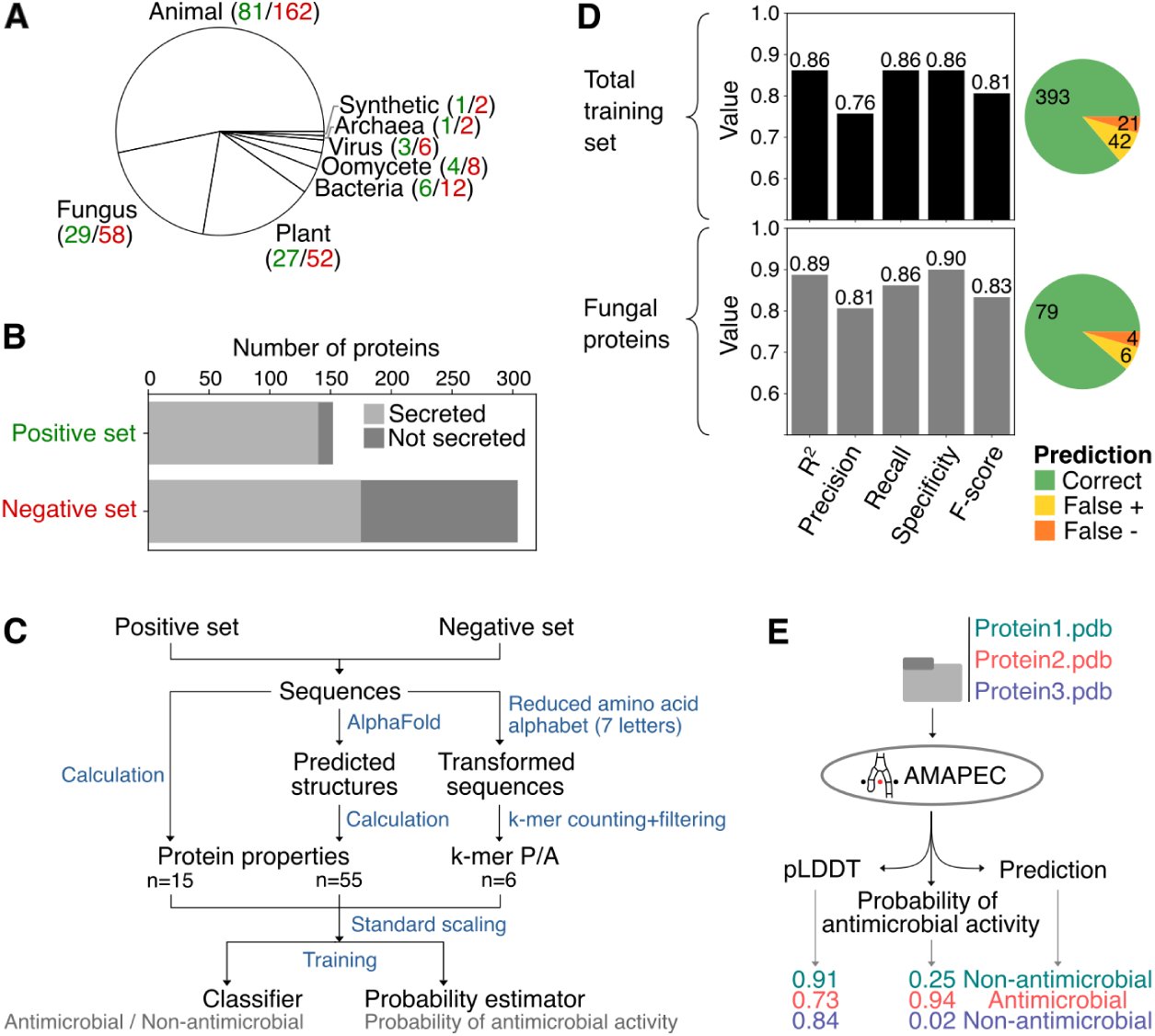
AMAPEC accurately predicts antimicrobial activities of fungal effector proteins. (**A**) Phylogenetic origin of the proteins in the dataset (green: number of proteins in positive dataset; red: number in negative dataset) used to train the predictor. (**B**) Number of proteins included in the training dataset and proportion of secreted proteins. (**C**) Diagram presenting a schematic overview of the training pipeline. In addition to physicochemical properties retrieved from protein sequences and structures, the pipeline considers the presence/absence (P/A) in protein sequences of six short motifs (k-mers) that are over- or underrepresented in the positive training set. These k-mers are encoded in a reduced amino acid alphabet, which groups amino acids according to their physicochemical properties. (n= number of variables). (**D**) Estimation of the classifier quality, based on “leave-one-out” cross-validation in the training dataset. The top bar plot and pie chart show quality estimates calculated on the total dataset (n=456), while the bottom charts analyze only the classifications of fungal proteins (n=87) during the “leave-one-out” cross validation. (**E**) Schematic overview of AMAPEC v1.0 showing its inputs and outputs with an example of three proteins.

We used AMAPEC to identify candidate antimicrobial proteins in the secretomes of three phylogenetically distant fungi with distinct lifestyles, namely the mycorrhizal glomeromycete *Rhizophagus irregularis*, the saprotrophic basidiomycete *Coprinopsis cinerea* and the ubiquitous plant-pathogenic ascomycete *Verticillium dahliae*, that causes vascular wilt disease in a wide diversity of host plants, including a variety of crops (*27*) (Fig. 2, A to C). Functional compositions of fungal secretomes remain largely elusive, as they comprise a significant proportion of proteins without functional annotations (Fig. 2, A) and proteins for which the assigned annotations are poorly informative (tables S5-S7). Surprisingly, AMAPEC prediction revealed that one third to half of each of their secretomes is composed of antimicrobial proteins (Fig. 2, B). Orthology analysis revealed that some antimicrobials are conserved across the three fungi, despite their wide phylogenetic spread, and thus ancestral to the three phyla (Fig. 2, C). Moreover, antimicrobials show significantly greater conservation across species (*i.e.* occurrence of orthologs) than non-antimicrobials (Fig. 2, C; Fisher’s exact test: odd’s ratio = 2.19, *P* = 3.6×10^-4^), suggesting that microbial antagonism predates other secretome functions.

**Fig. 2.**
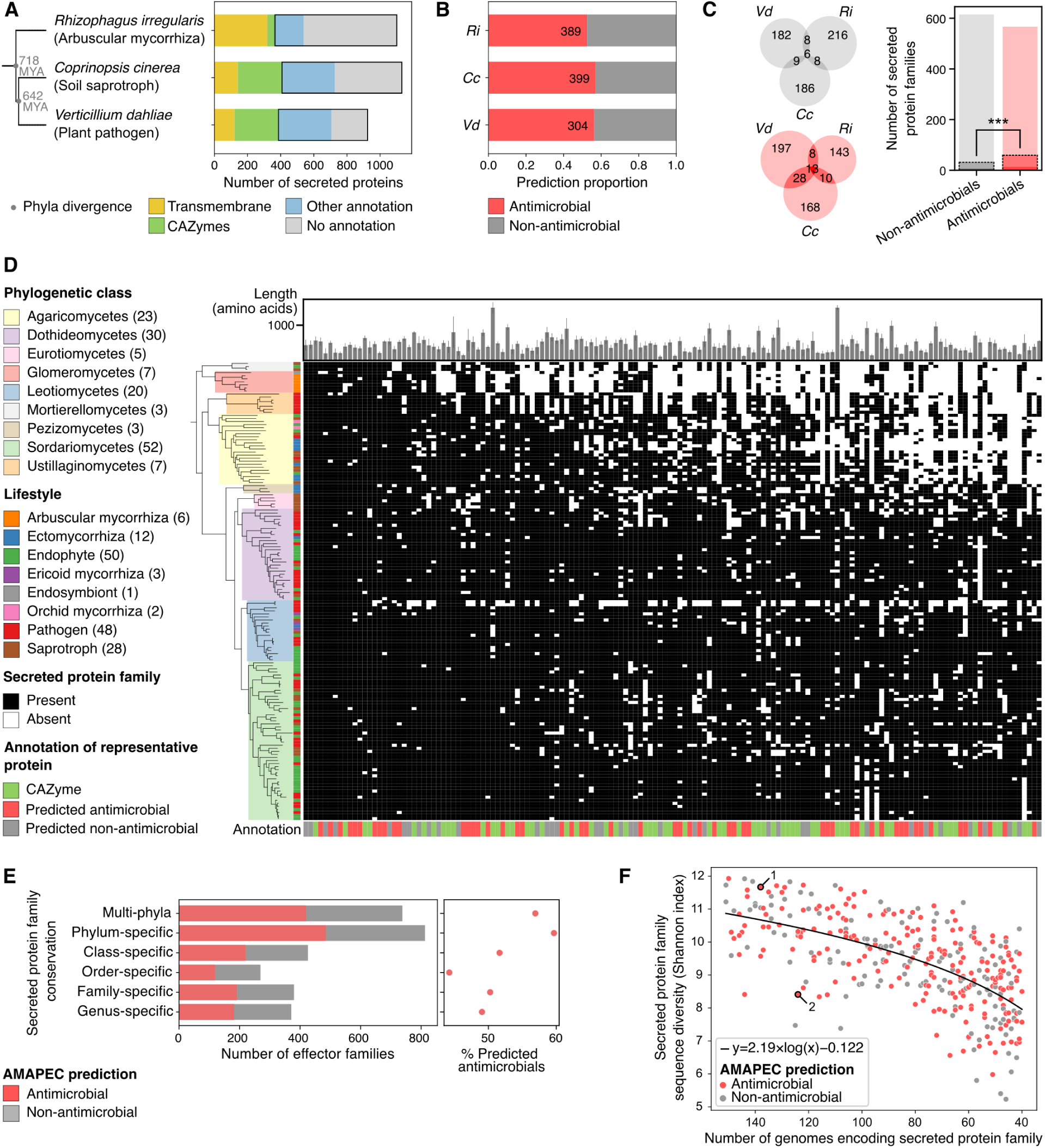
Predicted antimicrobials are abundant in fungal secretomes and exhibit high conservation. (**A**) Selected set of three fungi with broad phylogenetic diversity and distinct lifestyles, together with the functional annotation of their predicted secretomes. Parts of the secretomes framed with black rectangles were used for antimicrobial effector prediction with AMAPEC. The species-level phylogenetic tree was generated with phyloT (phylot.biobyte.de) based on NCBI taxonomy data (*50*). It is annotated with median taxa divergence times registered in the database TimeTree (*40*). (**B**) Results of AMAPEC antimicrobial activity prediction, highlighting proportions and numbers of predicted antimicrobials in the secretomes (excluding transmembrane proteins and CAZymes) of the arbuscular mycorrhizal Glomeromycota *Rhizophagus irregularis* (*Ri*), the saprophytic basidiomycete *Coprinopsis cinerea* (*Cc*) and the plant-pathogenic ascomycete *Verticillium dahliae* (*Vd*). (**C**) Orthology prediction analysis revealing the numbers of conserved predicted antimicrobial effectors (red Venn diagram) and non-antimicrobials (grey Venn diagram) across the three secretomes. On the right, a barplot shows numbers of secreted protein families and highlights with darker colors and a dash line the proportions of conserved antimicrobials and non-antimicrobials across the three secretomes. Fisher’s exact test (odd’s ratio = 2.19, *P* = 3.6×10^-4^) revealed an overrepresentation of conserved effectors among predicted antimicrobials. (**D**) Analysis of the 150 most conserved secreted protein families (according to orthology prediction) in a diverse dataset of 150 fungal genomes. On the left, a phylogeny describes the genomic dataset composition, with phylogenetic classes and lifestyles annotated in colors. In the central heatmap, each column corresponds to a secreted protein family with black and white squares highlighting the presence/absence of these families in each genome. Protein family conservation decreases from left to right. On top, a barplot shows average protein sequence lengths. At the bottom, colored bars highlight the CAZyme annotation or results of AMAPEC antimicrobial activity prediction performed on a representative member of each protein family. (**E**) On the left, a barplot depicts numbers of predicted antimicrobial and non-antimicrobial families (prediction performed on one representative protein per family, excluding families which representative protein was annotated as transmembrane protein or CAZyme) across effector family conservation levels in the 150-genome dataset. On the right, a scatterplot presents proportions of predicted antimicrobials along the same effector family conservation levels. These proportions follow a significantly descending trend according to Cochran-Armitage test (statistic = 5129.0, *P* < 2.36×10^-5^). (**F**) Intra-family sequence diversity indexes (Shannon alpha-diversity calculated from k-mer composition of amino acid sequences) of effector families is plotted as a function of decreasing family conservation across the dataset of 150 fungal genomes, with dot color corresponding to antimicrobial activity prediction on a representative member of each family. A logarithm function was fitted to these data. Two protein families labeled 1 and 2 include previously experimentally validated fungal antimicrobial proteins (*11*, *29*, *51*).

### Secreted antimicrobial proteins are conserved throughout the fungal kingdom

The evolution of antimicrobial proteins was further investigated in 150 genomes of diverse soil- and plant-associated fungi, spanning 3 phyla, 9 classes and 24 orders (fig. S4 and table S8). This dataset encompasses over 700 million years of fungal evolution, whereas pathogenicity towards land plants only evolved after plants colonized lands, about 500 million years ago. After classifying secreted proteins into sequence-related families through orthology prediction, antimicrobial activities were predicted for the most representative protein of each family (table S9). Remarkably, many of the most conserved protein families, which occur in fungi with diverse lifestyles, were predicted as antimicrobials (Fig. 2, D). We calculated the percentage of predicted antimicrobial families across different levels of conservation (Fig. 2, E) and identified a significant decrease from 56.9% of multi-phyla families to 49.1% of genus-specific families (Cochran-Armitage test for trend: statistic = 5129.0, *P* < 2.36×10^-5^). This overrepresentation of predicted antimicrobials among the most conserved secreted protein families signifies their ancient origins, preceding fungal phyla divergence, and corroborates that fungi have relied on antimicrobials long before establishing symbioses with multicellular eukaryotes such as land plants and animals (*28*). While the most conserved secreted protein families expectedly exhibit the most diverse sequences, certain predicted antimicrobial families, including previously characterized ribonuclease-like antimicrobials (*11*, *29*), display low sequence diversity, suggesting purifying selection (Fig. 2, F). Together, our results demonstrate that microbial antagonism through the secretion of antimicrobial proteins is an ancient and conserved trait that likely supports fungal fitness across a wide diversity of habitats.

### Effectors of plant-pathogenic fungi display antimicrobial activities

We recently showed that *V. dahliae* secretes effector proteins with selective antimicrobial activity to manipulate resident microbiota at various life stages, including host colonization (*7–9*, *14*). Intriguingly, we noticed that several *V. dahliae* effectors previously characterized to modulate host immunity have predicted antimicrobial properties (table S10), including VdCP1 (*30*), Vd424Y (also known as VdXyn4) (*31*, *32*) and Vd2LysM (*33*), a member of the LysM effector family which represses chitin-triggered plant immunity and has an ancient origin since family members occur throughout the fungal kingdom (*34*, *35*). Based on these findings, we hypothesized that plant-pathogenic fungi have evolved effectors to manipulate plant host physiology from ancestral antimicrobial proteins. In support of this hypothesis, we found that most functionally characterized effectors registered in PHI-base (*36*) (n=76/133), some of which are broadly conserved throughout our dataset of 150 genomes (Fig. 3, B), have predicted antimicrobial properties (Fig. 3, A and table S11). To validate these predictions, we subsequently selected five effectors for experimental validation: the LysM effector Ecp6 from the tomato leaf mould fungus *Cladosporium fulvum* (*37*), AGLIP1 from the root rot fungus *Rhizoctonia solani* (*38*), AVR-Pita from the rice blast fungus *Magnaporthe oryzae* (*39*) and Vd424Y (*31*, *32*) and VdCP1 (*30*) from *V. dahliae*. All proteins were heterologously produced in *Escherichia coli* and employed in antimicrobial activity assays using a diverse set of microbes including 12 bacteria, four yeasts and three filamentous fungi. Interestingly, despite their well-characterized host targets, all five effectors exhibited antimicrobial activities *in vitro* with highly distinct activity spectra at micromolar concentrations (Fig. 3, C and D, and fig. S6-S15). As homologs of all five proteins occur in fungi that do not live in association with plants, and most homologs in their protein families are predicted antimicrobials (fig. S5 and table S12-16), antimicrobial activity is likely ancestral and predates the evolution of host manipulation. Overall, our data suggest that pathogen effectors evolved from ancient antimicrobial proteins that retain their ancestral functions while acquiring the ability to manipulate host physiology.

**Fig. 3.**
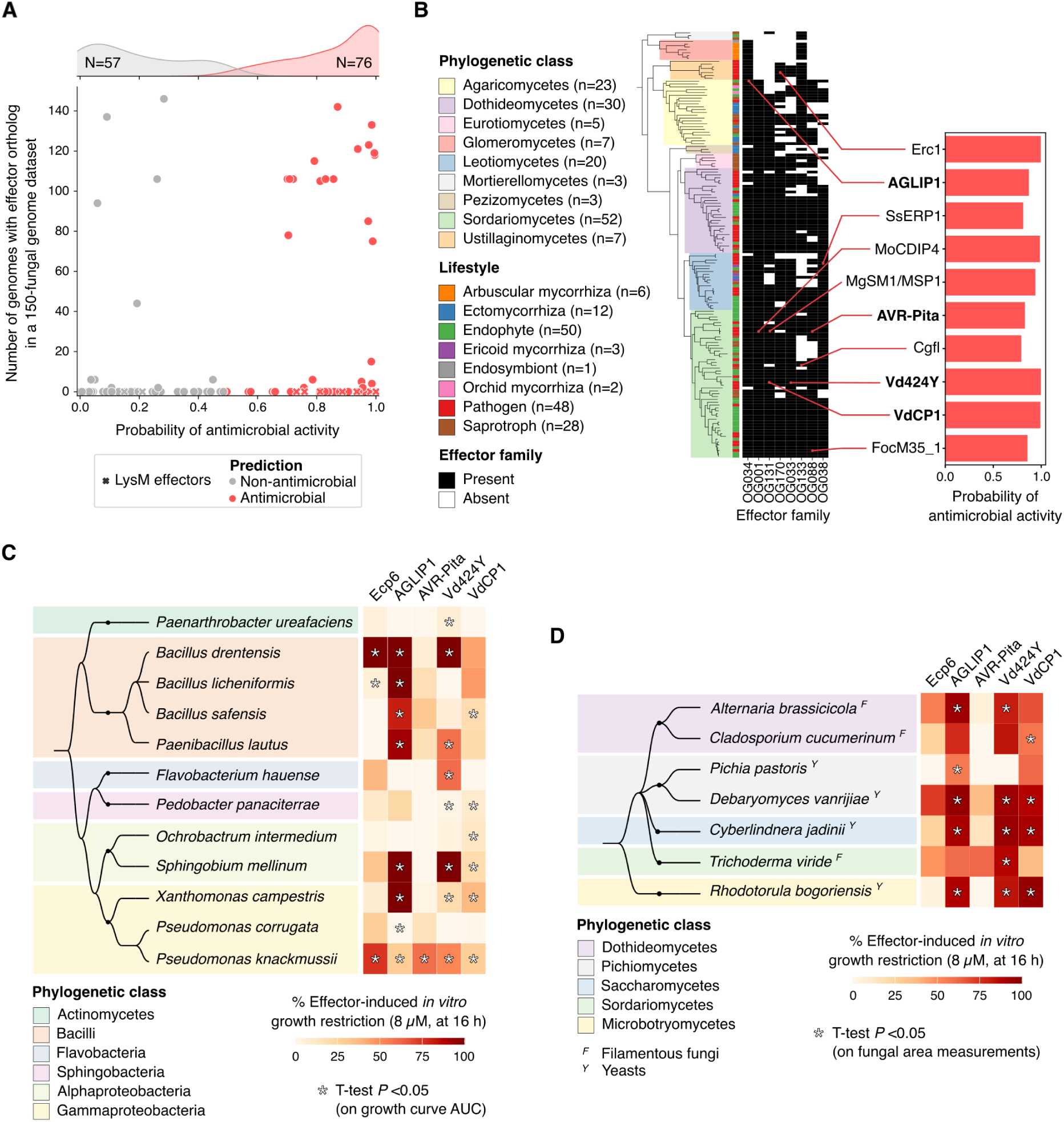
Immunomodulatory effectors also possess antimicrobial properties. (**A**) Scatterplot showing the antimicrobial activity prediction for fungal effectors from the curated PHI-base database (*36*) together with their number of orthologs in our dataset of 150 fungal genomes as a proxy for their conservation across the fungal kingdom. (**B**) Presence/absence patterns in the dataset of 150 fungal genomes of eight secreted protein families, including effectors that were previously demonstrated to act on plant hosts. On the right, a barplot shows the name of the previously studied effectors together with their AMAPEC-predicted antimicrobial activity. The eight families were selected because they include a highly similar protein to the PHI-base reference effectors (≥80% amino acid sequence identity), they show high conservation across the fungal kingdom (family represented in ≥100 genomes in the 150-genome dataset) and because the associated reference effectors are predicted antimicrobials with highly confident structure prediction (mean pLDDT ≥ 70). Effector names in bold indicate effectors selected for experimental validation of predicted antimicrobial activities. (**C**) Heatmap highlighting bacterial growth restriction induced by the presence of 8 µM of effector protein in the growth medium. A phylogenetically diverse set of 12 bacteria was used and is described with a species phylogeny on the left (generated with Taxallnomy (*52*)). Each heatmap column corresponds to a different fungal effector. Percentages of effector-induced growth restriction were calculated after 16 hours of growth. Asterisks highlight significant differences between from bacterial growth in presence and in absence of effector protein, identified with Student’s T-tests on area under curve (AUC) values. (**D**) Heatmap highlighting fungal growth restriction induced by the presence of 8 µM of effector protein in the growth medium. A phylogenetically diverse set of 7 fungi (4 yeasts and 3 filamentous fungi) was used and is described with species phylogeny on the left (tree generated with Taxallnomy (*52*)). Each heatmap column corresponds to a different fungal effector. Percentages of effector-induced growth restriction were calculated after 16 hours of growth. Asterisks highlight significant differences between from fungal growth in presence and in absence of effector protein, identified with Student’s T-tests. More details on the results of *in vitro* assays depicted on panels C and D can be found in fig. S6-S15.

### Dual roles of effectors in host manipulation and microbial antagonism

Since many fungal effectors have evolutionary conserved ancestral antimicrobial properties (Fig. 3), we hypothesize that, in addition to manipulation of host physiology, they antagonize microbial competitors during infection. To address this hypothesis, we focused on the *V. dahliae* effector Vd424Y, which is known to exert xylanolytic and cytotoxic activities *in planta* (*31, 32*). Vd424Y was previously demonstrated to localize into host cell chloroplasts and nuclei and, accordingly, carries a chloroplast transit peptide (cTP) and a nuclear localization signal (NLS). While the cTP is sparsely detected throughout the Vd424Y family (fig. S16, A) and shows a high degree of sequence variation (fig. S16, B), the occurrence of the NLS is restricted to a single clade that is exclusively composed of proteins secreted by plant pathogens and endophytes from the Sordariomycete genera *Verticillium*, *Fusarium* and *Cylindrocarpon* (Fig. 4, A), which diverged about 250 million years ago (*40*). This finding supports the hypothesis that contrary to antimicrobial properties, plant cell nuclear localization, which is required for immunomodulation and cytotoxicity *in planta* (*31*), is a trait that emerged recently in the family of Vd424Y homologs. To investigate whether Vd424Y functions in microbial competition, we first measured effector gene expression in a diverse set of ten soils, in absence of a plant host (fig. S17). We found that *V. dahliae* expresses the *Vd424Y* gene in these soils, suggesting functions beyond host physiology manipulation. Next, we tested if the antimicrobial activity of Vd424Y plays a role during plant colonization, by performing tomato inoculation experiments in a gnotobiotic system, allowing to study virulence contributions of effector genes in the presence and absence of host-associated microbiota (*41*). We identified a significant microbiota-dependent contribution of Vd424Y to *V. dahliae* virulence, as disease development was compromised upon *Vd424Y* deletion in the presence, but not in the absence of host-associated microbes (Fig. 4, B and C). This finding suggests that Vd424Y plays a role in microbiota manipulation during tomato plant infection, and thus that it retained its ancestral antimicrobial property throughout co-evolution with plant hosts.

**Fig. 4.**
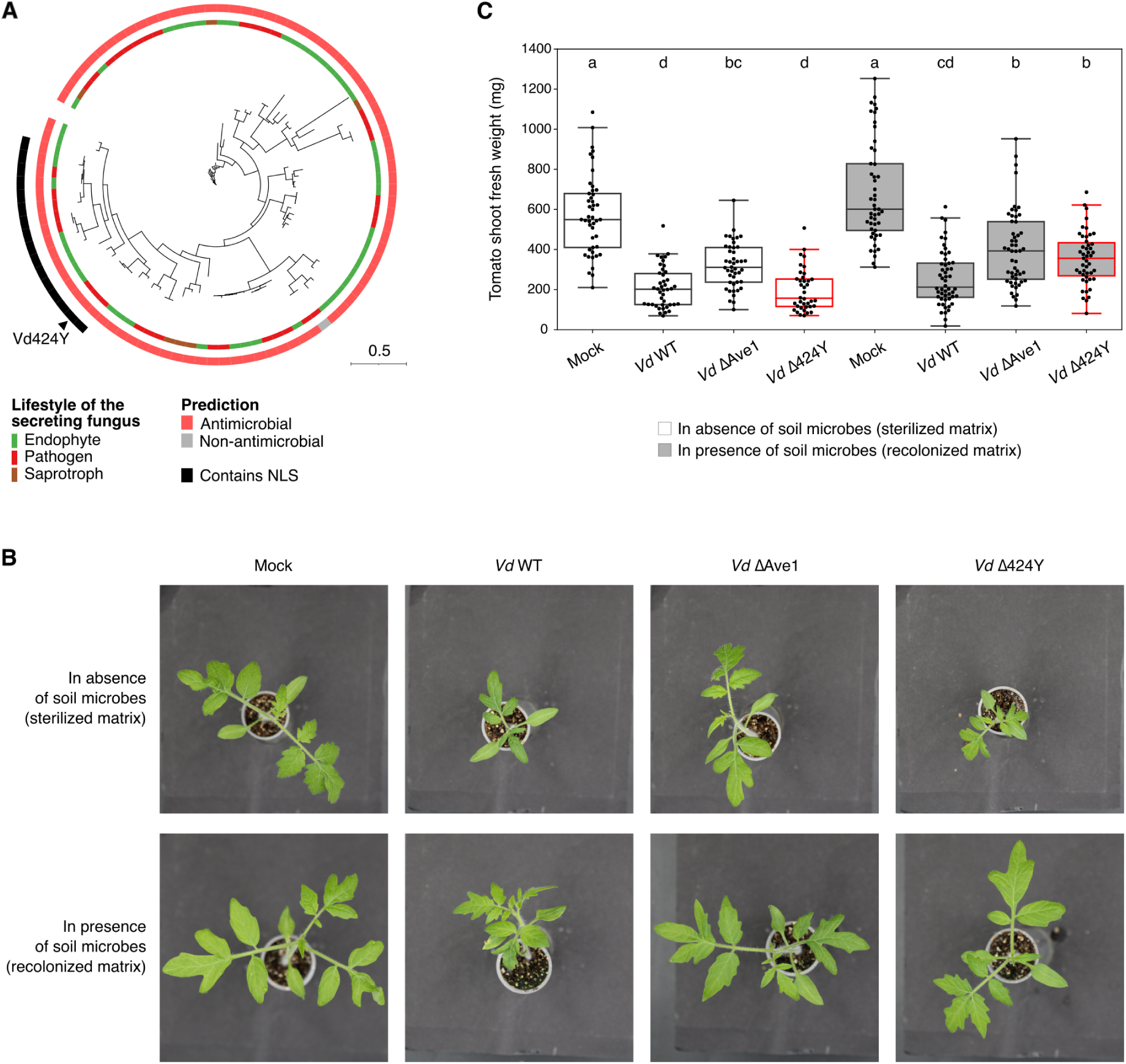
The effector Vd424Y acts in *Verticillium dahliae* antagonism towards plant microbiota members. (**A**) Maximum-likelihood phylogenetic tree (computed with IQ-TREE (*53*), model ‘LG’) of Vd424Y homologs in a dataset of 150 fungal genomes. These proteins occur in the same subfamily as Vd424Y (*i.e.* large clade identified in the total family phylogenetic tree, see fig. S5). The tree was manually rooted at the protein identified as the optimal outgroup by IQ-TREE and annotated with the lifestyles of the fungus secreting each protein, the results of antimicrobial activity as well as the occurrence of a nuclear localization signal (NLS), annotated using cNLS Mapper (*54*). The annotated NLS sequences in the effector family are all identical (fig. S16, B). (**B**) Representative photographs of tomato plants upon mock or fungal inoculation in a gnotobiotic system in absence (sterilized matrix; top) or presence (recolonized matrix; bottom) of a soil microbiota. Fungal strains used for plant infections include wild-type *V. dahliae* (*Vd* WT), *Vd424Y* deletion mutant (*Vd* Δ424Y), and as a control, an *Ave1* deletion mutant lacking the Ave1 effector previously demonstrated to contribute to *V. dahliae* virulence through both an antimicrobial activity targeting plant microbiota members and the manipulation of plant physiology (*7*, *41*). (**C**) Tomato shoot fresh weight values reflecting plant health after mock or fungal inoculation in a gnotobiotic system. Plants were grown axenically (sterilized matrix; white boxes) or in presence of microbes (recolonized matrix; grey boxes). The *Vd424Y* deletion mutant (*Vd* Δ424Y) is highlighted with red boxplot borders. Between 38 and 52 tomato plants were grown per condition, over three independent biological replicates. Letters on the boxplot highlight significant differences between treatments identified by ANOVA test (*P*<0.05) followed by Tukey HSD post-hoc test (adjusted *P*<0.05).

## Discussion

Before plants colonized land about 500 million years ago, fungi lived as saprotrophs and/or parasites of bacteria and primitive algae in (semi)aquatic environments (*42*). Pathogenicity towards land plants thus represents a relatively recent trait in the 1.5 billion year-long fungal evolutionary history, which evolved multiple times independently across the fungal tree of life (*43*). Effector-based strategies to manipulate host physiology are essential for the pathogenic lifestyle (*5*, *6*), yet their evolutionary origins remained elusive. The broad conservation of certain effector families throughout the fungal kingdom (Fig. 3, B), together with their occurrence in fungi that do not colonize plants, demonstrates ancient origins of these effectors and suggests their primary roles beyond plant host manipulation. As here we found that many effector proteins have ancestral antimicrobial properties, we identified a fundamental function of effectors in microbial antagonism, which likely supported fungal fitness in (semi)aquatic environments before fungi engaged in symbioses with land plants (*28*). In line with this discovery, previous studies identified peptides with dual functions in microbial antagonism and immunity modulation, suggesting these two protein activities may often co-occur given their functional complementarity (*44*, *45*). Moreover, a recent study identified that changes in protein domain organization may have repurposed an effector from antimicrobial to immunomodulator, thereby mediating a recent fungal lifestyle transition from saprotroph to plant symbiont (*46*). Hypothetically, the co-option of antimicrobial proteins for host manipulation mediated the compatibility of the first fungal symbioses with land plants and predates the evolution of a plethora of other molecular mechanisms that underly more specific pathogenic interactions. Indeed, some fungal effectors represent recent innovations, as evidenced by their lineage-specificity, and likely result from fungal adaptation to specific plant lineages (*15*, *47*). Such effectors are generally thought of as products of co-evolutionary “arms races” in which plant immune systems and fungi aim to detect and overcome detection, respectively (*1*, *3*). While this study focuses on the occurrence of antimicrobial properties among the most conserved effector families, it paves the way for further research on effector evolution, particularly to investigate how recently evolved antimicrobial effectors may mediate host and niche adaptation (*48*). More generally, this study, and more particularly the AMAPEC software, will support the discovery of novel fungal antimicrobial proteins and assist studies on their modes-of-action, which currently remain largely unknown. Furthermore, the broad variety of fungal secreted antimicrobials and their crucial role in supporting host colonization are important factors to consider when designing novel biocontrol strategies to protect crops efficiently from pathogenic fungi. Finally, the evolutionary trajectories identified in this study are likely to be relevant beyond plant colonization, given the need for human- and animal-pathogenic fungi to manipulate immune responses of their hosts during infection too (*49*).

## Materials and Methods

### Curation of a training dataset for antimicrobial activity prediction

To develop the AMAPEC predictor, a positive training set of antimicrobial proteins (table S1) was curated from literature. Only proteins for which antimicrobial activity has been experimentally demonstrated *in vitro* (*i.e.* restricting the growth of bacteria and/or fungi in culture medium) were selected. While not restraining the dataset to proteins encoded by any phylogenetic group, we paid particular attention to include all the fungal antimicrobial proteins reported in scientific literature. Importantly, secretion signal peptides were removed from sequences (SignalP v6.0 (*55*)), since the antimicrobial function of proteins occurs after secretion. Considering sequence lengths of fungal secreted proteins, peptide with mature sequence lengths below 40 amino acids were excluded (fig. S1), not to enrich the protein set in AMPs, for which dedicated predictors exist (*19*, *56–60*). By largely spanning the size range of typical effector proteins, this protein set should support the prediction of effector antimicrobial activity without bias towards the recognition of short AMPs, that are the most described antimicrobial proteins in the literature.

A negative training dataset was assembled by gathering presumable non-antimicrobial proteins (table S2). As previously suggested (*19*, *57*), this negative set was curated by retrieving proteins which functional annotation does not suggest any antimicrobial activity from the UniProt database (*61*). To do so, Gene Ontology (GO) terms associated to antimicrobial activity were filtered out (*i.e.* GO:0090729, GO:0001878, GO:0045087, GO:0050830, GO:0050829, GO:0042742, GO:0071222, GO:0071224, GO:0001530, GO:0031640, GO:0050832). Additionally, only well-annotated proteins without any known function in microbial antagonism or immunity were selected. To prevent strong effects of potential misjudgment during the curation process, the negative training dataset includes twice as many non-antimicrobial proteins as there are antimicrobials in the positive set. For each antimicrobial in the positive set, two presumably non-antimicrobial proteins encoded by the same organism (or a close relative) and with similar sizes (+/− 4 amino acids) were included in the negative set. We paid attention to include at least as many secreted proteins (signal peptide detected and removed with SignalP (*55*)) in the negative set as in the positive set, not to bias the prediction towards apoplastically released proteins. Finally, since 11 proteins in the positive training set were annotated or described as ribonucleases, 11 ribonucleases, unlikely to exert antimicrobial functions according to their annotation (for instance, involved in transfer RNA maturation) were included in the negative set.

### Calculation of protein properties

The AMAPEC predictor was trained on a set of 70 numerical variables reflecting protein physicochemical properties (table S3). Some of these values (n=15) were calculated from amino acid sequences, using R v4.2.0 and the library Peptides v2.4.4 (*62*). However, to better describe the physicochemistry of proteins, their predicted structures were used to calculate 55 numerical values per protein that reflect structural properties. Protein structures were predicted using AlphaFold v2.0 (*63*) with parameters “--max_template_date=2021-05-14 –preset=casp14”. Structure properties were calculated from AlphaFold best models (“ranked_0.pdb” output files) using Python v3.11.5 and the PDB parser implemented in Biopython v1.78 (*64*). Some previously published code and formula from diverse sources (*65–67*) (details in table S3) were implemented in AMAPEC’s Python scripts. For properties linked to protein secondary structures, DSSP v3.0.0 (*68*, *69*) was used to assign individual amino acids to different types of secondary structures. Additionally, pocket structures in proteins were predicted using Fpocket v4.0.2 (*70*), and information related to their number, size and properties were implemented as variables.

In addition to sequence- and structure-derived physicochemical properties, the presence/absence of certain k-mers in the protein sequences was implemented as variable. To reduce sequence complexity, a reduced amino acid alphabet was used, as previously implemented in various machine learning methods applied to protein sequences (*57*, *71*). A novel 7-letter alphabet based on amino acid properties was designed, to define k-mers that may represent key motifs in protein physicochemistry: small amino acids (G, A) were mapped to ‘0’, nucleophilic amino acids (S, T, C) were mapped to ‘1’, hydrophobic amino acids (V, L, I, M, P) were mapped to ‘2’, aromatic amino acids (F, Y, W) were mapped to ‘3’, acidic amino acids (D, E) were mapped to ‘4’, amide amino acids (N, Q) were mapped to ‘5’ and basic amino acids (H, K, R) were mapped to ‘6’. The k-mer compositions of transformed sequences in the training set was profiled using MerCat2 v1.0 (*72*), with k-mer sizes 3, 4, 5 and 6 amino acids. Then, chi-squared multiple testing was computed, as implemented in function feature_selection.SelectFdr(chi2, alpha=0.05) of Python library scikit-learn v1.2.1 (*73*), to identify k-mers that are over- or under-represented in antimicrobial protein sequences. This aimed to select for k-mers that are likely biologically meaningful, and to prevent later overfitting of our prediction model that can occur if training is performed on numerous variables which combination describes protein sequences in too much detail. With chi-squared testing, six k-mers (five 4-mers and one 3-mer) of interest were identified. Their presence/absence was implemented in the set of protein properties used to train the AMAPEC predictor (table S3).

### Classifier training and quality estimation

Numerical variables reflecting properties of our 456 proteins were standardized using function preprocessing.StandardScaler() of Python library scikit-learn v1.2.1 (*73*). Then, a Support Vector Machines (SVM) classifier with a linear kernel was trained using function svm.SVC() from scikit-learn. The correct the imbalance of the training set (152 proteins in the positive set and 304 in the negative set), the weight of antimicrobials was set to 2 and the weight of non-antimicrobials to 1. A second model was trained to predict the probability of antimicrobial activity, by computing Platt scaling over the SVM binary classifier. To do so, the function calibration.CalibratedClassifierCV(method=’sigmoid’, cv=’prefit’) from sci-kit learn was used. Both models were exported using function dump() from Python library joblib v1.2.0.

Due to the small size of the training dataset (n=456), classifier quality testing was performed through leave-one-out cross-validation. As implemented in function cross_val_score(cv=KFold(n_splits=456)) of scikit-learn, 456 SVM classifiers were trained with a train/test split of 455/1 to classify individual proteins using as a basis, protein properties in the rest of the dataset. Protein classifications into “antimicrobial” or “non-antimicrobial” were then analyzed by counting numbers of true positives, false positives, true negatives and false negatives. These counts allowed the estimation of the overall classifier accuracy (R^2^) but also its precision, recall, specificity and F-score. Such quality estimates were also calculated by exclusively taking the classification correctness of fungal proteins into account, to identify if the predictor is suited for the annotation of fungal proteins. Finally, the predictor was tested on seven recently characterized fungal proteins demonstrated to have antimicrobial activities (*22–26*), after structure prediction with ESMFold v1.0.3 (*74*) (table S4).

A bash pipeline allowing both the calculation of protein properties and antimicrobial activity prediction using the trained predictors was written, resulting in the software AMAPEC v1.0 (https://github.com/fantin-mesny/amapec), (developed and tested on operating system GNU/Linux Ubuntu v20.04.3 LTS).

### Secretome analysis on three phylogenetically distant fungi

Sets of proteins associated to the published genomes of three phylogenetically distant fungi with distinct lifestyles were downloaded: *Verticillium dahliae* JR2 (*75*) (annotation *VDAG_JR2 v.4.0* downloaded from the database Ensembl Fungi (*76*)), *Coprinopsis cinerea* AmutBmut pab1-1 (*77*) (annotation *Copci_AmutBmut1 v1.0* downloaded from the database JGI Mycocosm (*78*)) and *Rhizophagus irregularis* DAOM197198 (*79*). SignalP v6.0 (*55*) was then used to predict secretion signal peptides in protein sequences and thereby define the secretomes of these fungi. Sequences with removed signal peptides were used in all subsequent analyses. Functional annotation of proteins in these secretomes was carried out using emapper v2.0 (*80*) and the database EggNog v5 (*81*). CAZymes and transmembrane proteins were specifically annotated in these secretomes using dbCAN v4.0 and TMBed v1.0.0 respectively (*82*, *83*). Structure predictions were computed with ESMFold v1.0.3 (*74*), using default parameters, of all secreted proteins besides CAZymes. The structures of two proteins from *V. dahliae* (VDAG_JR2_Chr4g10970 and VDAG_JR2_Chr1g22375) could not be predicted due to high computational requirements linked to their size (>2500 amino acids) and were excluded from our analyses. To validate that the quality of protein structures predicted by ESMFold is sufficient, structure prediction for 626/635 non-CAZyme secreted proteins of *V. dahliae* was also performed with AlphaFold v2.0 (*63*) with parameters --max_template_date=2021-05-14 --preset=casp14, with nine predictions failing due to high computational requirements. Average structure pLDDT values were compared between AlphaFold and ESMFold (fig. S18). Then, AMAPEC v1.0 was used to predict the antimicrobial activity of proteins secreted by *V. dahliae*, *C. cinerea* and *R. irregularis*, while excluding CAZymes and transmembrane proteins. ESMFold (*74*)-predicted structures, which pLDDT confidence scores are in tables S5-7, were used as an input. To analyze the conservation of predicted antimicrobials and non-antimicrobials, an orthology prediction was computed on the three fungal secretomes with OrthoFinder v2.5.5 (*84*). The conservation of antimicrobials and non-antimicrobials was studied independently, after subsetting the OrthoFinder-generated ‘orthogroups’ tables to only contain proteins from each group.

### Comparative genomics in a dataset of 150 fungal genomes

A set of 150 fungal genomes was assembled, based on a previously published dataset of 120 fungal genomes for which fungal lifestyles had been manually curated (*85*). This dataset was supplemented with 23 genomes of plant-pathogenic fungi and 7 genomes of Glomeromycetes (see details in table S8). An orthology prediction was performed on total sets of annotated proteins with OrthoFinder v2.5.5 (*84*) to generate a phylogenomic tree with the implemented method ‘STAG’ (*86*). This genome-scale phylogeny is displayed on Fig. 2D and fig. S4. In all 150 genomes, sets of proteins carrying SignalP v6.0 (*55*)-annotated signal peptides were considered to form secretomes. A second orthology prediction with OrthoFinder v2.5.5 was performed on these 150 secretomes. In each secreted protein family (*i.e.* ‘orthogroup’) defined by this orthology prediction, the most central and representative protein was identified with phylorep v0.1 (*87*) using OrthoFinder-generated gene trees (relying on method FastTree (*88*)) as inputs. CAZymes and transmembrane proteins among these family representatives were annotated using dbCAN v4.0 and TMBed v.1.0.0, respectively (*82*, *83*). The structures of other representative proteins were predicted with ESMFold v1.0.3 (*74*) then submitted to antimicrobial activity prediction with AMAPEC v1.0. Protein family conservation analyses were performed considering the presence/absence of families in each genome (*i.e.* considering the occurrence of orthologs but not the numbers of paralogs). Specifically, each family was assigned a conservation level according to the clade (phylum, class, order, family or genus) its occurrence is restricted to, in the 150 genome dataset. Proportions of protein families which representative members are predicted antimicrobials/non-antimicrobials were analyzed, excluding CAZyme- and transmembrane protein-encoding families. Then, sequence diversity within the 600 most conserved secreted protein families (excluding CAZymes) was estimated through k-mer based Shannon index calculation with MerCat2 v1.0 (*72*) using a k-mer size of 3 amino acids.

### Analysis of predicted antimicrobials in PHI-base

Sequences of proteins that have been studied for their contribution to host-pathogen interactions were downloaded from the PHI-base database (*36*) (accessed in January 2024). To identify proteins previously studied for their contribution to fungal virulence in the *V. dahliae* JR2 secretome, SignalP v6.0 (*55*)-predicted secreted proteins in the *V. dahliae* genome were blasted against the downloaded PHI-base sequences using blastp v2.5.0 with parameters -evalue 0.0001 - max_target_seqs 1 and additional filtering to only keep hits with more than 95% sequence identity to query proteins (table S10). To investigate more generally if previously characterized fungal effectors have antimicrobial activities, the PHI-base dataset was subsetted to only retain proteins from fungi that have secretion signal peptides according to SignalP v6.0 (*55*) and that were classified as “effectors (plant avirulence determinants)” in the database. After structure prediction with ESMFold v1.0.3 (*74*), the antimicrobial activity of these proteins was predicted with AMAPEC v1.0. To estimate the conservation of these effectors across the fungal kingdom, the protein family of the most similar protein in the 150-genome dataset was considered and the number of genomes in which it occurs was used as a proxy for effector conservation (Fig. 3, A). To identify previously characterized fungal effectors with confidently predicted antimicrobial activity and broad conservation across the fungal kingdom (Fig. 3, B), the following filtering criteria were used: highly confident effector structure prediction with an average pLDDT>70, predicted antimicrobial activity, presence of an effector homolog with more than 80% sequence identity in the 150-genome dataset and representation of the associated family in more than 100/150 genomes.

### Effector protein production and purification

For heterologous production of the AGLIP1, AVR-Pita, Ecp6, Vd424Y and VdCP1 effectors, protein sequences were retrieved from PHI-base (*36*), then codon-optimized nucleotide sequences encoding for mature proteins were subcloned into pET-15b (AVR-Pita, Vd424Y, AGLIP1), pET-28a(+) (VdCP1) or pETSUMO (Ecp6) expression vectors. All the constructs with an N-terminal His_6_ tag (Gene Universal, Newark, DE, USA) were transformed into *Escherichia coli* BL21 or Shuffle cells by heat shock (45 s at 42 °C, 5 min on ice). The transformed BL21 cells (AGLIP1, AVR-Pita, Vd424Y and VdCP1) were grown at 37 °C with constant shaking at 180 rpm in 2x YT medium (16 g/l tryptone, 10 g/l yeast extract, 5 g/l NaCl) containing 100 μg/ml ampicillin. Protein production was induced with 1 mM isopropyl-ß-d-thiogalactoside (IPTG) when cultures reached an optical density (OD_600_) of 2.0. Induction was performed for 2 h at 37 °C for AVR-Pita and Vd424Y, 4 h at 37 °C for AGLIP1, and 2 h at 42 °C for VdCP1, all with constant shaking at 180 rpm. For protein extraction, the bacterial cells were pelleted and then resuspended in denaturing 6 M guanidinium chloride (GdmCl), 10 mM β-mercaptoethanol and 10 mM Tris, pH 8.0, and incubated overnight at 4 °C with continuous rotation. The lysate was centrifuged at 16,000 × *g* for 1 h and the resulting cleared supernatant was collected for protein purification by metal affinity chromatography using a nickel His60 Ni Superflow Resin (Takara, San Jose, CA, USA) column in the ÄKTA go™ protein purification system (Cytiva, Marlborough, MA, USA). After column equilibration with 6 M GdmCl, 10 mM Tris at pH 8.0, denatured protein was loaded onto the column and weakly bound protein was washed off with 6 M GdmCl, 10 mM Tris, 20 mM Imidazole at pH 8.0 before the His_6_-tagged protein was eluted with 6 M GdmCl, 10 mM Tris, 200 mM Imidazole at pH 8.0. Purified proteins were dialyzed (Spectra/Por™ 3 RC Dialysis Membrane Tubing 3500 Dalton molecular weight cut-off; Spectrum Laboratories – Rancho Dominguez, CA, USA) sequentially against 20 volumes of buffers with decreasing GdmCl concentrations (table S17) to promote refolding. Each dialysis step lasted a minimum of 24 h. Finally, depending on their isoelectric point, the proteins were dialyzed twice against 200 volumes of 30 mM sodium phosphate buffer (pH 7.0 for Vd424Y, pH 5.8 for AGLIP1) or 30 mM potassium phosphate buffer (pH 7.5 for AVR-Pita, pH 6.5 for VdCP1). Final concentrations were determined with a Qubit 4 Fluorometer (Invitrogen – Waltham, MA, USA). For the heterologous protein production of Ecp6 ( *89*, *90*), *E. coli* Shuffle cells carrying pETSUMO-Ecp6 were grown at 37°C in lysogeny broth (LB) containing 50 μg/mL kanamycin until an OD_600_ of 0.8. Protein expression was induced with 0.2 mM IPTG, followed by incubating at 18°C overnight. Cells were pelleted and resuspended in cell lysis buffer (50 mM Tris-HCl pH 7.5, 150 mM NaCl, 10% glycerol, 6 mg/ml lysozyme, 2 mg/ml deoxycholic acid, 60 µg/ml DNase I (Roche – Basel, Switzerland; ref. 04536282001) and one protease inhibitor cocktail pill (Roche; ref. 11836170001)), then incubated either at room temperature for 3 h or at 4 °C overnight with stirring. The lysate was centrifuged at 20,000 × *g* for 1 h, and the cleared supernatant was collected for protein purification. His_6_-tagged SUMO-Ecp6 was purified using His60 Ni Superflow Resin, pre-equilibrated with 20 mM Tris, 150 mM NaCl, 5 mM imidazole, pH 8.0. Bound protein was eluted with 20 mM Tris, 150 mM NaCl, 300 mM imidazole, pH 8.0. The His_6_-SUMO affinity tag was removed by incubation overnight with ULP-1 protease (Sigma-Aldrich – St. Louis, MO, USA; ref. SAE0067) in dialysis buffer (20 mM Tris, 100 mM NaCl, 2% glycerol, pH 8.0). Non-cleaved fusion protein was removed by a second round of affinity purification using His60 Ni Superflow resin, and the flow-through containing cleaved Ecp6 was dialyzed overnight. The final protein concentration was determined spectrophotometrically at 280 nm.

### In vitro microbial growth inhibition assays

A diverse set of twelve bacteria previously isolated from tomato plants (*41*) were grown on LB agar plates at room temperature in the dark, then transferred into low-salt tryptic soy broth (ls-TSB; 17 g/l tryptone, 3 g/l soy peptone, 0.5 g/l sodium chloride, 2.5 g/l dipotassium phosphate, and 2.5 g/l glucose) and grown overnight at 28 °C while shaking at 180 rpm. Overnight cultures were resuspended to the final OD_600_ = 0.025 in equal parts of fresh ls-TSB and AVR-Pita, Vd424Y, AGLIP1, VdCP1 or Ecp6 in phosphate buffer of the corresponding pH (see above; final protein concentration: 8 µM) or in the respective phosphate buffer only as a control. Total volumes of 100 µl were incubated in clear 96-well flat-bottom polystyrene tissue culture plates in a CLARIOstar plate reader (BMG Labtech – Ortenberg, Germany) at 25 °C with double orbital shaking every 15 minutes (10 seconds at 300 rpm). The optical density was measured every 15 minutes at 600 nm. After OD_600_ normalization, areas under curves in presence and absence of effector proteins were calculated using the trapezoidal method.

Four yeasts, i.e. *Pichia pastoris* GS115, *Cyberlindnera jadinii* (DSM 70167), *Debaryomyces vanrijiae* (DSM 70252) and *Rhodotorula bogoriensis* (DSM 70872), were grown on potato dextrose agar (PDA; ROTH – Karlsruhe, Germany, ref. X931) at room temperature in the dark then transferred into 0.05X potato dextrose broth (PDB) and grown overnight at 25 °C while shaking at 180 rpm. Overnight cultures were resuspended to the final OD_600_ = 0.025 in equal parts of fresh 0.05X PDB and AVR-Pita, Vd424Y or AGLIP1 in phosphate buffer of the corresponding pH (see above; final protein concentration: 8 µM) or in the phosphate buffer only as a control. Additionally, spores from filamentous fungal strains of *Alternaria brassicicola*, *Cladosporium cucumerinum*, *Trichoderma viride* (from our in-house culture collection) were harvested from PDA plates after culture at room temperature in the dark, then separated from the mycelium with a sterile 40 μm nylon filter (VWR – Radnor, PA, USA) and suspended in equal parts of 0.05X PDB and AVR-Pita, Vd424Y or AGLIP1 in phosphate buffer of the corresponding pH (see above; final protein concentration: 8 µM) or in the respective phosphate buffer only as a control to a final concentration of 10^4^ spores/ml. Total volumes of 100 µl were incubated in clear 96-well flat-bottom polystyrene tissue culture plates at 25 °C overnight. For both filamentous fungi and yeasts, fungal growth was imaged using an CKX41 inverted microscope (Olympus – Shinjuku City, Japan) with DP20 camera (Olympus). Images were analyzed with ImageJ (*91*): Each image was first subjected to binarization and next to particle analysis to measure total particle area.

### Analysis of the evolutionary histories of effectors with validated antimicrobial activities

The evolutionary histories of effectors AGLIP1, AVR-Pita, Vd424Y and VdCP1 were analyzed by reconstructing the protein family phylogenies. All sequences in the protein families (defined through orthology prediction in the 150-genome dataset) including these effectors were used to reconstruct maximum-likelihood phylogenetic trees using IQ-TREE v2.0.3 (model ‘LG’, default settings) (*53*) after multiple sequence alignment with MAFFT v7.310 (default parameters) (*92*). Since fungal LysM effectors are thought to represent a single effector family but exhibit large sequence diversity and therefore occur in multiple protein families defined by orthology prediction, the LysM effector family was identified through functional annotation, following a previously introduced procedure (*34*). In the 150-fungal genome dataset, and additionally in the genome of the fungus *Cladosporium fulvum* which secretes the well-characterized Ecp6 LysM effector (*37*), functional domains were annotated using InterProScan v5.65-97.0 (*93*). Secreted proteins containing LysM domains (IPR036779, IPR018392, IPR045030), but not any other functional domains (*e.g.* chitinase domains), were considered as LysM effectors. All annotated LysM effectors were used to reconstruct a protein phylogenetic tree with IQ-TREE (model ‘LG’) after sequence alignment with MAFFT, as performed for the four other effector families. In all five effector families, antimicrobial activities were predicted using AMAPEC v1.0 after structure prediction with ESMFold v1.0.3 (*74*). Protein family phylogenetic trees were visualized and annotated with antimicrobial activity prediction results using iTOL (*94*).

The recent evolution of the Vd424Y effector family was further studied focusing on the subfamily (clade identified on the total protein family tree, fig. S5) containing Vd424Y. As previously (*32*), chloroplastic transit peptides and nuclear localization signals were annotated using ChloroP v1.1 (*95*) and cNLS Mapper (*54*) (used online with default parameters in May 2025: https://nls-mapper.iab.keio.ac.jp/), respectively. To analyze sequence variations in the Vd424Y subfamily, subfamily-members were aligned using MAFFT and a figure was generated with iTOL (*94*), referring to a previously published domain annotation of Vd424Y to highlight the domain organization of the proteins (*32*).

### Measurements of *V. dahliae* effector gene expression in soils

Soil extracts were prepared from 10 previously sampled soils with distinct properties and microbial communities (*96*). To do so, 20 g of soil (stored at 4 °C) were mixed with 100 ml of sterile water (1:5, w/v) followed by incubation at room temperature for 2 days. Soil particles were removed by centrifugation at 4,000 × *g* for 30 min, and the resulting supernatant was used as soil extract.

Conidiospores of *V. dahliae* JR2 were harvested from mycelium cultured on PDA plates for 7 days. The spores were washed once with sterile water and collected by centrifugation at 10,000 × *g* for 2 min. Spore concentration was determined by counting with a hemocytometer, and 1×10⁶ spores were inoculated into 10 ml of PDB in 50 ml flasks. Cultures were incubated at 22 °C with shaking at 130 rpm for 2 days. Mycelia were then collected, rinsed with sterile water, and transferred into 10 ml of prepared soil extracts in new 50 ml flasks. After 2 days of incubation, the mycelia were collected using Miracloth, rinsed with sterile water, and blotted dry with tissue paper. The samples were transferred to 2 ml tubes containing two 2.3 mm iron beads, flash-frozen in liquid nitrogen, and ground using a tissue lyser. Total RNA was extracted using TRIzol reagent (Thermo Fisher Scientific – Waltham, MA, USA). For each sample, 500 ng of RNA was reverse transcribed using the HiScript® III RT SuperMix for qPCR (+gDNA Wiper) (Vazyme Biotech Co., Ltd. - Nanjing, China; ref. R323-01). The resulting cDNA was diluted 10 fold with nuclease-free water. Quantitative real-time PCR was performed using SsoAdvanced™ Universal SYBR® Green Supermix (Bio-Rad, Hercules, CA, USA). Primers 5’-TGTTACCAAAGCAGCACACAAGG-3’ and 5’-CCTTATGCCTCGTTCCCTTCCAC-3’ were used to amplify the Ave1-encoding gene (positive control, (*7*)), primers 5’-GCAAGCGAGGACTGACAAGATC-3’ and 5’-CGACGGAATGGACGGCGTG-3’ were used to amplify the Tom1-enconding gene (negative control, (*97*)), newly designed primers 5’-TCGGGCGGTTTCTACTACTC-3’ and 5’-TGTTGTTCTTCCAGCTGACG-3’ were used to amplify the Vd424Y-encoding gene (VDAG_JR2_Chr5g00880), and the primers 5’-CGAGTCCACTGGTGTCTTCA-3’ and 5’-CCTCAACGATGGTGAACTT-3’ were used to amplify the glyceraldehyde 3-phosphate dehydrogenase gene (GAPDH, house-keeping gene). The PCR cycling conditions were as follows: initial denaturation at 95 °C for 3 mins, followed by 40 cycles of denaturation at 95 °C for 10 s and annealing/extension at 60 °C for 30 s with fluorescence signal collection. After amplification, a melt curve analysis was performed from 65 °C to 95 °C, increasing by 0.5 °C every 5 seconds, to verify the specificity of the PCR products. Gene expression levels of effector genes were normalized to the *V. dahliae* GAPDH using the ΔCt method.

### Vd424Y gene deletion in Verticillium dahliae

Gene deletion was performed following previously published protocols for *V. dahliae* protoplast transformation (*98*) and CRISPR-Cas9-based fungal genome editing (*99*). After culturing on PDA plates at room temperature for 7-15 days in the dark, 3×10^7^ spores of *V. dahliae* JR2 were harvested from the mycelium surface and inoculated into 100 ml of liquid Complete Medium (CM) composed of 0.6% yeast extract (Duchefa – Haarlem, The Netherlands; ref. Y1333), 0.6% casein hydrolysate (ROTH; ref. AE41) and 1% sucrose (VWR; ref. 0335) in milli-Q water. After 20 hours in culture at 28 °C with agitation, mycelium was harvested on a Falcon® 40μm nylon filter (ref. 352340) and washed with a sterile 0.7 M NaCl solution. Then, the harvested mycelium was incubated in 10 ml of sterile-filtered driselase solution (0.2% of enzyme in 0.7 M NaCl, Sigma-Aldrich – St. Louis, MO, USA; ref. D9515) for 2.5 hours at 33 °C with agitation. The resulting solution was passed through a 40 μm nylon filter, then centrifugated at 3000 × *g* for 5min. After supernatant removal, the protoplast pellet was resuspended in 1 ml of sterile STC (20% Sucrose, 10 mM Tris-HCl pH 8.0 and 50 mM CaCl_2_ in milli-Q water) and centrifuged at 3000 × *g* for 5 min. This last step was repeated twice to thoroughly wash the protoplasts. Protoplasts were counted under a microscope, and their concentration was adapted to 5 × 10^6^ protoplasts/ml in STC. After adding 1% of DMSO, protoplast solutions were kept at −80 °C.

To delete the gene of interest, two single guide RNA (sgRNA) were designed using CRISPick (https://portals.broadinstitute.org/gppx/crispick/public) to target the upstream and downstream regions of the Vd424Y-encoding gene (VDAG_JR2_Chr5g00880): CATACGTCCTGTTCAGCCGG (upstream) and GCCATCCGACCAGCATTCAG (downstream). A blastn was performed (with parameter --word_size 11) to check that these sgRNA protospacer sequences only occur near the targeted gene in the *V. dahliae* JR2 genome. Oligonucleotides with sequences corresponding to the designed sgRNA protospacer in between sequences 5’-AAGCTAATACGACTCACTATA-3’ and 5’-GTTTTAGAGCTAGAAATAGCAAG-3’ were ordered. To synthesize sgRNA from these oligonucleotides, 1 µl of oligonucleotide (100 µM) was mixed with 1 µl of oligonucleotide 5’-AAAAGCACCGACTCGGTGCCACTTTTTCAAGTTGATAACGGACTAGCCTTATTTTAA CTTGCTATTTCTAGCTCTAAAAC-3’ (100 µM) and 8 µl of nuclease-free water. This mixture was incubated in a thermocycler for annealing with the following program: 95 °C for 5 min, 95 °C to 85 °C at 2 °C/sec, 85 °C to 25 °C at −0.1 °C/sec. Then, 2.5 µl of dNTPs (10 mM each), 2 µl of NEBuffer™ r2.1 (10X, New England Biolabs – Ipswich, MA, USA; ref. B6002S), 0.5 µl T4 DNA polymerase (New England Biolabs; ref. M0203S) and 5 µl nuclease-free water were added to the mixture, followed by an incubation at 12 °C for 20 min. Products of this reaction were purified with a Monarch^®^ PCR & DNA Cleanup Kit (New England Biolabs; ref. T1030L). Concentration in DNA was then measured with a NanoDrop device. To transcribe the DNA molecules into sgRNA, 2 µg of DNA was mixed with 6µl of RNA NTPs (25 mM each), 1.5 µl of T7 buffer (10X), 1.5µl of HiScribe^®^ T7 polymerase (New England Biolabs; ref. E2040S), 1 µl of DTT (0.1 M) and nuclease-free water up to 20 µl total volume. After incubation overnight at 37 °C, 14 µl of nuclease-free water, 4 µl of RQ1 DNase buffer (10X, Promega – Walldorf, Germany; ref. M198A) and 2 µl of RQ1 DNAse (Promega; ref. M610A) were added in the reaction tube. An incubation of 30 min at 37 °C followed to digest remaining DNA. sgRNA molecules were then purified using RNA Clean & Concentrator (Zymo Research – Irvine, CA, USA; ref. R1017 & R1018) kit. Purified sgRNA were stored at −80 °C until transformation.

Double-stranded donor DNA corresponding to the 50 bp-sequence upstream of the expected Cas9-mediated cut in the fungal genome, followed by the 50 bp-sequence downstream of the second expected cut was synthesized. Through homologous recombination, this donor DNA promoted genome repair excluding our target gene upon double-stranded cuts by Cas9.

The commercial enzyme EnGen® Spy Cas9 HF1 (New England Biolabs; ref. M0667M) was used for protoplast transformation. First, 4 µM of this Cas9 enzyme was mixed to 2X of NEBuffer™ r3.1 (New England Biolabs; ref. B6003S) and 0.3 µg/µl of sgRNA. The mixture was incubated at 25 °C for 30 min to bind sgRNA and enzyme. Both prebound Cas9-sgRNA complexes were then mixed with 200 µl of fungal protoplasts (5 × 10^6^ protoplasts/ml), 20 pmol of double-stranded donor DNA and 6 µg of telomeric vector pTEL-Hyg (*99*) containing a hygromycin resistance gene. After 30min of incubation on ice, 1.5 ml of PEG-STC (60% polyethylene glycol 4000, 20% of sucrose, 10 mM of Tris-HCl pH 8.0 and 50 mM CaCl_2_ in milli-Q water) was added to the mixture and gently mixed by tube rotation. This tube was incubated at room temperature for 15 min. Then, 5 ml of TB3 medium (3% yeast extract, 3% casein hydrolysate and 20% sucrose in water) were added to the mixture, stimulating protoplast regeneration over an 18 hour-incubation in the dark at room temperature.

Regeneration medium was centrifuged at 3000 × *g* for 5 min. The supernatant was removed and pelleted fungi were resuspended in sterile water to be plated on PDA medium containing 50 µg/ml of hygromycin. Plates were incubated at room temperature for 5 days. Then, visible colonies were screened by PCR using a pair of primers targeting ∼500 bp upstream and downstream of the gene of interest (forward: 5’-ACATATCGCGACGAGTTCCC-3’, reverse: 5’-CTCTTCTTCTCGAGCGACCC-3’). One colony for which the amplicon size was the one expected upon successful gene deletion was identified. Sanger sequencing of the amplicon confirmed the successful gene deletion. After cultivation of this mutant on PDA+hygromycin plate, spores were harvested and inoculated on hygromycin-free PDA medium. A colony that successfully grew in absence of hygromycin and that was confirmed to have lost the pTEL-Hyg vector was used as Vd424Y deletion line (Δ424Y).

Before to be used in plant recolonization experiments, growth of the newly generated mutant line was tested to verify it is not impaired in growth. *V. dahliae* JR2 wild-type and the Δ424Y mutant were cultured on PDA plates for 10 days. Conidiospores were harvested and washed twice with sterile milli-Q water. The final concentration was adjusted to 1 × 10⁶ spores/ml in 1 ml of PDB medium. This spore solution was incubated horizontally at 25 °C with shaking for 48 hours. Following incubation, fungal cells were collected by centrifugation at 13,000 rpm for 5 minutes, and 900 µl of the supernatant was carefully removed to avoid disturbing the pellet. As an internal control, 1 ng of synthetic spike-in plasmid (*100*) was added to each sample, and genomic DNA was extracted from the resulting pellets using the DNeasy PowerSoil Pro Kit (Qiagen – Hilden, Germany; ref. 47014), following the manufacturer’s protocol. *V. dahliae* biomass was quantified by real-time PCR using species-specific primers VdITS1-F (5’-AAAGTTTTAATGGTTCGCTAAGA-3’) and STVe1-R (5’-CTTGGTCATTTAGAGGAAGTAA-3’) targeting the internal transcribed spacer (ITS) region. The spike-in plasmid was amplified in the same solutions using primers qRT-Spike-F (5’-TTTCTTTTCCAAGGTTTGTGC-3’) and qRT-Spike-R (5’-AACATTTACCCTGCTTGTAGCTCT-3’). A fungal growth index was calculated from both amplifications’ Ct values: index = 2^-(CtITS/CtSpike)^. The growth index values of *V. dahliae* wild-type and Δ424Y mutant were then compared to check that the Δ424Y mutant line is not impaired in growth (fig. S19).

### Tomato plant inoculation assays in a gnotobiotic system

The protocol applied here follows an adaptation of the FlowPot gnotobiotic system (*101*) to tomato plant inoculations with *V. dahliae* strains (*41*). FlowPot substrate, a 1:1 mixture of peat and vermiculite (Balster Einheitserde – Frödenberg, Germany; LIMERA Gartenbauservice – Geldern-Walbeck, Germany) was sterilized by two consecutive autoclaving rounds. Following substrate sterilization, FlowPot units were created by filling truncated 50 ml Luer lock syringes (Terumo Europe, Leuven, Belgium) with sterilized substrate and autoclaving for a third time on a liquid cycle. Additionally, a recolonized condition was created, by mixing sterilized substrate with non-sterile substrate in a 9:1 ratio, followed by an incubation at room temperature overnight. To remove toxic compounds, that may accumulate during autoclaving, the substrate was flushed using 30 ml of sterile milliQ-water using a vacuum system. Further, substrate was enriched using 30 ml of half-strength Murashige & Skoog medium (Duchefa; ref. M0222). Following FlowPot preparation, tomato seeds (*Solanum lycopersium* L.; cultivar “MoneyMaker”) were surface sterilized as described previously (*102*), stratified at 8 ℃ for 24 hours and sown into each FlowPot unit. Subsequently, up to 5 FlowPot units were placed into Microbox containers with 4 air filters (SacO2; Deinze, Belgium) and kept in a greenhouse (17 hours of light at 23 ℃ followed by 7 hours of darkness at 22 ℃). After 14 days of growth, tomato plants were inoculated using *Verticillium dahliae* wild-type and Δ424Y strains as well as ΔAve1 (*7*) a control. To this end, Microboxes were opened in a sterile hood, and plants were carefully uprooted from the substrate. Roots were rinsed using sterile milliQ-water and subsequently placed into a *V. dahliae* spore suspension, containing 10^6^ spores/ml. After 8 minutes of incubation, plants were placed back into the FlowPots and Microboxes were placed back into the greenhouse chamber. Symptom development was assessed at 14 days-post inoculation by measuring tomato shoot fresh weight.

### Statistics

Fisher’s exact tests were computed using the function *stats.fisher_exact* in SciPy v1.13.0 (*103*). Mann-Whitney U tests were performed using the function *stats.mannwhitneyu* of SciPy v1.13.0. Cochrane-Armitage tests for trend were calculated with the function *stats.contingency_tables.Table.test_ordinal_association* of statsmodels v0.14.0 (*104*). In case of multiple testing, p-values from the tests mentioned above were adjusted using Benjamini-Hochberg correction with the function *stats.multitest.multipletests(method=’fdr_bh’)* of statsmodels v0.14.0. Measurements from *in vitro* microbial growth restriction assays were analyzed in R v4.4.1, first by using the *shapiro.test* function and Q-Q plots to assess normality of the datasets (Shapiro-Wilk test *P*>0.05) then since all data was normally distributed, pairwise one-sided Student’s t-tests comparing microbial growth in presence and absence of protein were performed using the function *t.test(alternative=’less’)*. To analyze the tomato shoot fresh weight measurements, a square-root transformation (in R: *lm(sqrt(ShootFreshWeight)∼Condition)*) was applied to reach normal distribution of the data (Shapiro-Wilk test *P*>0.05), then statistical comparison between treatments was performed using an ANOVA test and a post-hoc Tukey HSD test (functions *aov* and *TukeyHSD* of R v4.2.0). Letters reflecting the significant differences between treatments were obtained with function *multcompLetters4* from R package multcompView v0.1-1 (*105*).

## Code availability

Data and scripts used for analyses in this study are available or linked at https://github.com/fantin-mesny/Scripts_analysis_ancient_fungal_antimicrobials.

## Acknowledgments

This work was supported by the Deutsche Forschungsgemeinschaft (DFG, German Research Foundation), through the funding of F.M.’s Walter Benjamin position (Project ID: ME 6064/1-1, Project number: 508411006). Y.S. acknowledges funding through an Overseas Research Fellowship from the Japan Society for the Promotion of Science while ALM acknowledges receipt of a postdoctoral research fellowship funded by the ‘Fundación Ramón Areces’. B.P.H.J.T. acknowledges funding by the Alexander von Humboldt Foundation in the framework of an Alexander von Humboldt Professorship endowed by the German Federal Ministry of Education and Research and is furthermore supported by DFG under Germany’s Excellence Strategy – EXC 2048/1 – Project ID: 390686111, by the DFG – Project ID 458090666 / CRC1535/1, and Project ID: FOR 5682, and additionally acknowledges support from iHEAD (NRW Profilbildung ID: PB22-025A). We thank Michael F. Seidl for critical feedback.

## References

1. D. E. Cook, C. H. Mesarich, B. P. H. J. Thomma, Understanding plant immunity as a surveillance system to detect invasion. Annu. Rev. Phytopathol. 53, 541–563 (2015).

2. G. Z. Han, Origin and evolution of the plant immune system. New Phytol. 222, 70–83 (2019).

3. J. D. G. Jones, B. J. Staskawicz, J. L. Dangl, The plant immune system: From discovery to deployment. Cell 187, 2095–2116 (2024).

4. V. Müller, R. J. de Boer, S. Bonhoeffer, E. Szathmáry, An evolutionary perspective on the systems of adaptive immunity. Biol. Rev. 93, 505–528 (2018).

5. I. Stergiopoulos, P. J. G. M. De Wit, Fungal effector proteins. Annu. Rev. Phytopathol. 47, 233–263 (2009).

6. G. Doehlemann, B. Ökmen, W. Zhu, A. Sharon, Plant pathogenic fungi. Microbiol. Spectr. 5 (2017).

7. N. C. Snelders, H. Rovenich, G. C. Petti, M. Rocafort, G. C. M. van den Berg, J. A. Vorholt, J. R. Mesters, M. F. Seidl, R. Nijland, B. P. H. J. Thomma, Microbiome manipulation by a soil-borne fungal plant pathogen using effector proteins. Nat. Plants 6, 1365–1374 (2020).

8. N. C. Snelders, G. C. Petti, G. C. M. van den Berg, M. F. Seidl, B. P. H. J. Thomma, An ancient antimicrobial protein co-opted by a fungal plant pathogen for in planta mycobiome manipulation. Proc. Natl. Acad. Sci. 118, e2110968118 (2021).

9. N. C. Snelders, J. C. Boshoven, Y. Song, N. Schmitz, G. L. Fiorin, H. Rovenich, G. C. M. van den Berg, D. E. Torres, G. C. Petti, Z. Prockl, L. Faino, M. F. Seidl, B. P. H. J. Thomma, A highly polymorphic effector protein promotes fungal virulence through suppression of plant-associated Actinobacteria. New Phytol. 237, 944–958 (2023).

10. E. A. Chavarro-Carrero, N. C. Snelders, D. E. Torres, A. Kraege, A. López-Moral, G. C. Petti, W. Punt, J. Wieneke, R. García-Velasco, C. J. López-Herrera, M. F. Seidl, B. P. H. J. Thomma, The soil-borne white root rot pathogen *Rosellinia necatrix* expresses antimicrobial proteins during host colonization. PLOS Pathog. 20, e1011866 (2024).

11. B. Ökmen, P. Katzy, L. Huang, R. Wemhöner, G. Doehlemann, A conserved extracellular Ribo1 with broad-spectrum cytotoxic activity enables smut fungi to compete with host-associated bacteria. New Phytol. 240, 1976–1989 (2023).

12. D. Gómez-Pérez, M. Schmid, V. Chaudhry, Y. Hu, A. Velic, B. Maček, J. Ruhe, A. Kemen, E. Kemen, Proteins released into the plant apoplast by the obligate parasitic protist *Albugo* selectively repress phyllosphere-associated bacteria. New Phytol. 239, 2320–2334 (2023).

13. H. X. Chang, Z. A. Noel, M. I. Chilvers, A β-lactamase gene of *Fusarium oxysporum* alters the rhizosphere microbiota of soybean. Plant J. 106, 1588–1604 (2021).

14. A. Kraege, W. Punt, A. Doddi, J. Zhu, N. Schmitz, N. C. Snelders, B. P. H. J. Thomma, Undermining the cry for help: The phytopathogenic fungus Verticillium dahliae secretes an antimicrobial effector protein to undermine host recruitment of antagonistic Pseudomonas bacteria. bioRxiv, 2025.06.09.658588 (2025).

15. M. Möller, E. H. Stukenbrock, Evolution and genome architecture in fungal plant pathogens. Nat. Rev. Microbiol. 15, 756–771 (2017).

16. L. J. Ma, H. C. Van Der Does, K. A. Borkovich, J. J. Coleman, M. J. Daboussi, A. Di Pietro, M. Dufresne, M. Freitag, M. Grabherr, B. Henrissat, P. M. Houterman, S. Kang, W. B. Shim, C. Woloshuk, X. Xie, J. R. Xu, J. Antoniw, S. E. Baker, B. H. Bluhm, A. Breakspear, D. W. Brown, R. A. E. Butchko, S. Chapman, R. Coulson, P. M. Coutinho, E. G. J. Danchin, A. Diener, L. R. Gale, D. M. Gardiner, S. Goff, K. E. Hammond-Kosack, K. Hilburn, A. Hua-Van, W. Jonkers, K. Kazan, C. D. Kodira, M. Koehrsen, L. Kumar, Y. H. Lee, L. Li, J. M. Manners, D. Miranda-Saavedra, M. Mukherjee, G. Park, J. Park, S. Y. Park, R. H. Proctor, A. Regev, M. C. Ruiz-Roldan, D. Sain, S. Sakthikumar, S. Sykes, D. C. Schwartz, B. G. Turgeon, I. Wapinski, O. Yoder, S. Young, Q. Zeng, S. Zhou, J. Galagan, C. A. Cuomo, H. C. Kistler, M. Rep, Comparative genomics reveals mobile pathogenicity chromosomes in *Fusarium*. Nature 464, 367–373 (2010).

17. R. de Jonge, M. D. Bolton, A. Kombrink, G. C. M. Van Den Berg, K. A. Yadeta, B. P. H. J. Thomma, Extensive chromosomal reshuffling drives evolution of virulence in an asexual pathogen. Genome Res. 23, 1271–1282 (2013).

18. Y. Sato, R. Bex, G. C. M. van den Berg, P. Santhanam, M. Höfte, M. F. Seidl, B. P. H. J. Thomma, Starship giant transposons dominate plastic genomic regions in a fungal plant pathogen and drive virulence evolution. Nat. Commun. 16, 1–17 (2025).

19. M. Torrent, D. Andreu, V. M. Nogués, E. Boix, Connecting peptide physicochemical and antimicrobial properties by a rational prediction model. PLoS One 6, e16968 (2011).

20. G. Wang, The antimicrobial peptide database is 20 years old: Recent developments and future directions. Protein Sci. 32, e4778 (2023).

21. F. Wan, F. Wong, J. J. Collins, C. de la Fuente-Nunez, Machine learning for antimicrobial peptide identification and design. Nat. Rev. Bioeng. 2, 392–407 (2024).

22. R. Eichfeld, L. K. Mahdi, C. De Quattro, L. Armbruster, A. B. Endeshaw, S. Miyauchi, M. J. Hellmann, S. Cord-Landwehr, D. Peterson, V. Singan, K. Lail, E. Savage, V. Ng, I. V. Grigoriev, G. Langen, B. M. Moerschbacher, A. Zuccaro, Transcriptomics reveal a mechanism of niche defense: two beneficial root endophytes deploy an antimicrobial GH18-CBM5 chitinase to protect their hosts. New Phytol. 244 (2024).

23. F. Chen, L. Ou, H. Wu, L. Huang, Y.-P. Chen, Expression and characterization of the antifungal protein PtAFP from *Pyrenophora tritci-repentis* by synonymous codon bias in *Escherichia coli*. Proc. SPIE 13208, Third Int. Conf. Biomed. Intell. Syst. (IC-BIS 2024) 13208, 13–19 (2024).

24. K. de Guillen, L. Mammri, J. Gracy, A. Padilla, P. Barthe, F. Hoh, M. Lahfa, J. Rouffet, Y. Petit-Houdenot, T. Kroj, M.-H. Lebrun, Zymoseptoria tritici effectors structurally related to killer proteins UmV-KP4 and UmV-KP6 are toxic to fungi, and define extended protein families in fungi. bioRxiv, 2024.10.14.618152 (2024).

25. Z. Sorger, P. Sengupta, K. Beier-Heuchert, J. Bautor, J. E. Parker, E. Kemen, G. Doehlemann, GH25 lysozyme mediates tripartite interkingdom interactions and microbial competition on the plant leaf surface. bioRxiv, 2025.04.04.647216 (2025).

26. L. Florez, V. M. Flores-Núñez, C. S. Francisco, E. Holtgrewe Stukenbrock, The fungal effector AvrStb6 regulates the wheat pathobiome. Zenodo, doi: 10.5281/ZENODO.15852925 (2025).

27. E. F. Fradin, B. P. H. J. Thomma, Physiology and molecular aspects of *Verticillium* wilt diseases caused by *V. dahliae* and *V. albo-atrum*. Mol. Plant Pathol. 7, 71–86 (2006).

28. N. C. Snelders, H. Rovenich, B. P. H. J. Thomma, Microbiota manipulation through the secretion of effector proteins is fundamental to the wealth of lifestyles in the fungal kingdom. FEMS Microbiol. Rev. 46, fuac022 (2022).

29. G. J. Kettles, C. Bayon, C. A. Sparks, G. Canning, K. Kanyuka, J. J. Rudd, Characterization of an antimicrobial and phytotoxic ribonuclease secreted by the fungal wheat pathogen *Zymoseptoria tritici*. New Phytol. 217, 320–331 (2018).

30. Y. Zhang, Y. Gao, Y. Liang, Y. Dong, X. Yang, J. Yuan, D. Qiu, The *Verticillium dahliae* SnodProt1-like protein VdCP1 contributes to virulence and triggers the plant immune system. Front. Plant Sci. 8, 289292 (2017).

31. L. Liu, Z. Wang, J. Li, Y. Wang, J. Yuan, J. Zhan, P. Wang, Y. Lin, F. Li, X. Ge, *Verticillium dahliae* secreted protein Vd424Y is required for full virulence, targets the nucleus of plant cells, and induces cell death. Mol. Plant Pathol. 22, 1109–1120 (2021).

32. D. Wang, J. Y. Chen, J. Song, J. J. Li, S. J. Klosterman, R. Li, Z. Q. Kong, K. V. Subbarao, X. F. Dai, D. D. Zhang, Cytotoxic function of xylanase VdXyn4 in the plant vascular wilt pathogen *Verticillium dahliae*. Plant Physiol. 187, 409–429 (2021).

33. A. Kombrink, H. Rovenich, X. Shi-Kunne, E. Rojas-Padilla, G. C. M. van den Berg, E. Domazakis, R. de Jonge, D. J. Valkenburg, A. Sánchez-Vallet, M. F. Seidl, B. P. H. J. Thomma, *Verticillium dahliae* LysM effectors differentially contribute to virulence on plant hosts. Mol. Plant Pathol. 18, 596–608 (2017).

34. R. de Jonge, B. P. H. J. Thomma, Fungal LysM effectors: extinguishers of host immunity? Trends Microbiol. 17, 151–157 (2009).

35. A. Kombrink, B. P. H. J. Thomma, LysM effectors: Secreted proteins supporting fungal life. PLOS Pathog. 9, e1003769 (2013).

36. M. Urban, A. Cuzick, J. Seager, V. Wood, K. Rutherford, S. Y. Venkatesh, J. Sahu, S. Vijaylakshmi Iyer, L. Khamari, N. De Silva, M. C. Martinez, H. Pedro, A. D. Yates, K. E. Hammond-Kosack, PHI-base in 2022: A multi-species phenotype database for pathogen– host Interactions. Nucleic Acids Res. 50, D837–D847 (2022).

37. R. de Jonge, H. P. Van Esse, A. Kombrink, T. Shinya, Y. Desaki, R. Bours, S. Van Der Krol, N. Shibuya, M. H. A. J. Joosten, B. P. H. J. Thomma, Conserved fungal LysM effector Ecp6 prevents chitin-triggered immunity in plants. Science (80-. ). 329, 953–955 (2010).

38. S. Li, X. Peng, Y. Wang, K. Hua, F. Xing, Y. Zheng, W. Liu, W. Sun, S. Wei, The effector AGLIP1 in *Rhizoctonia solani* AG1 IA triggers cell death in plants and promotes disease development through inhibiting PAMP-triggered immunity in *Arabidopsis thaliana*. Front. Microbiol. 10, 483550 (2019).

39. G. Xiao, N. Laksanavilat, S. Cesari, K. Lambou, M. Baudin, A. Jalilian, M. J. Telebanco-Yanoria, V. Chalvon, I. Meusnier, E. Fournier, D. Tharreau, B. Zhou, J. Wu, T. Kroj, The unconventional resistance protein PTR recognizes the *Magnaporthe oryzae* effector AVR-Pita in an allele-specific manner. Nat. Plants 10, 994–1004 (2024).

40. S. Kumar, G. Stecher, M. Suleski, S. B. Hedges, TimeTree: a resource for timelines, timetrees, and divergence times. Mol. Biol. Evol. 34, 1812–1819 (2017).

41. W. Punt, J. Park, H. Roevenich, A. Kraege, N. Schmitz, J. Wieneke, N. C. Snelders, G. L. Fiorin, A. López-Moral, E. A. Chavarro-Carrero, G. C. Petti, K. Wippel, B. P. H. J. Thomma, A gnotobiotic system reveals multifunctional effector roles in plant-fungal pathogen dynamics. bioRxiv, 2025.03.27.645772 (2025).

42. T. Y. James, F. Kauff, C. L. Schoch, P. B. Matheny, V. Hofstetter, C. J. Cox, G. Celio, C. Gueidan, E. Fraker, J. Miadlikowska, H. T. Lumbsch, A. Rauhut, V. Reeb, A. E. Arnold, A. Amtoft, J. E. Stajich, K. Hosaka, G. H. Sung, D. Johnson, B. O’Rourke, M. Crockett, M. Binder, J. M. Curtis, J. C. Slot, Z. Wang, A. W. Wilson, A. Schüßler, J. E. Longcore, K. O’Donnell, S. Mozley-Standridge, D. Porter, P. M. Letcher, M. J. Powell, J. W. Taylor, M. M. White, G. W. Griffith, D. R. Davies, R. A. Humber, J. B. Morton, J. Sugiyama, A. Y. Rossman, J. D. Rogers, D. H. Pfister, D. Hewitt, K. Hansen, S. Hambleton, R. A. Shoemaker, J. Kohlmeyer, B. Volkmann-Kohlmeyer, R. A. Spotts, M. Serdani, P. W. Crous, K. W. Hughes, K. Matsuura, E. Langer, G. Langer, W. A. Untereiner, R. Lücking, A. Büdel, D. M. Geiser, A. Aptroot, P. Diederich, I. Schmitt, M. Schultz, R. Yahr, D. S. Hibbett, F. Lutzoni, D. J. McLaughlin, J. W. Spatafora, R. Vilgalys, Reconstructing the early evolution of Fungi using a six-gene phylogeny. Nature 443, 818–822 (2006).

43. M. A. Guerreiro, E. H. Stukenbrock, Fungal plant pathogens. Curr. Biol. 35, R480–R484 (2025).

44. C. Y. Huang, K. Araujo, J. N. Sánchez, G. Kund, J. Trumble, C. Roper, K. E. Godfrey, H. Jin, A stable antimicrobial peptide with dual functions of treating and preventing citrus Huanglongbing. Proc. Natl. Acad. Sci. U. S. A. 118, e2019628118 (2021).

45. D. Wu, L. Fu, W. Wen, N. Dong, The dual antimicrobial and immunomodulatory roles of host defense peptides and their applications in animal production. J. Anim. Sci. Biotechnol. 13, 141 (2022).

46. R. Eichfeld, A. B. Endeshaw, M. J. Hellmann, B. M. Moerschbacher, A. Zuccaro, Domain gain or loss in fungal chitinases drives ecological specialization toward antagonism or immune suppression. bioRxiv, 2025.06.16.659886 (2025).

47. P. van Dam, L. Fokkens, S. M. Schmidt, J. H. J. Linmans, H. Corby Kistler, L. J. Ma, M. Rep, Effector profiles distinguish formae speciales of *Fusarium oxysporum*. Environ. Microbiol. 18, 4087–4102 (2016).

48. F. Mesny, M. Bauer, J. Zhu, B. P. H. J. Thomma, Meddling with the microbiota: Fungal tricks to infect plant hosts. Curr. Opin. Plant Biol. 82, 102622 (2024).

49. A. C. Sexton, B. J. Howlett, Parallels in fungal pathogenesis on plant and animal hosts. Eukaryot. Cell 5, 1941–1949 (2006).

50. C. L. Schoch, S. Ciufo, M. Domrachev, C. L. Hotton, S. Kannan, R. Khovanskaya, D. Leipe, R. McVeigh, K. O’Neill, B. Robbertse, S. Sharma, V. Soussov, J. P. Sullivan, L. Sun, S. Turner, I. Karsch-Mizrachi, NCBI Taxonomy: a comprehensive update on curation, resources and tools. Database 2020 (2020).

51. N. Istifadah, J. A. Saleeba, P. A. McGee, Isolates of endophytic *Chaetomium* spp. inhibit the fungal pathogen *Pyrenophora tritici-repentis* in vitro. Can. J. Bot. 84, 1148–1155 (2006).

52. T. Sakamoto, J. M. Ortega, Taxallnomy: an extension of NCBI Taxonomy that produces a hierarchically complete taxonomic tree. BMC Bioinformatics 22, 1–23 (2021).

53. B. Q. Minh, H. A. Schmidt, O. Chernomor, D. Schrempf, M. D. Woodhams, A. Von Haeseler, R. Lanfear, E. Teeling, IQ-TREE 2: New models and efficient methods for phylogenetic inference in the genomic era. Mol. Biol. Evol. 37, 1530–1534 (2020).

54. S. Kosugi, M. Hasebe, M. Tomita, H. Yanagawa, Systematic identification of cell cycle-dependent yeast nucleocytoplasmic shuttling proteins by prediction of composite motifs. Proc. Natl. Acad. Sci. U. S. A. 106, 10171–10176 (2009).

55. F. Teufel, J. J. Almagro Armenteros, A. R. Johansen, M. H. Gíslason, S. I. Pihl, K. D. Tsirigos, O. Winther, S. Brunak, G. von Heijne, H. Nielsen, SignalP 6.0 predicts all five types of signal peptides using protein language models. Nat. Biotechnol. 40, 1023–1025 (2022).

56. P. K. Meher, T. K. Sahu, V. Saini, A. R. Rao, Predicting antimicrobial peptides with improved accuracy by incorporating the compositional, physico-chemical and structural features into Chou’s general PseAAC. Sci. Rep. 7, 1–12 (2017).

57. D. Veltri, U. Kamath, A. Shehu, Deep learning improves antimicrobial peptide recognition. Bioinformatics 34, 2740–2747 (2018).

58. T.-T. Lin, L.-Y. Yang, I.-H. Lu, W.-C. Cheng, Z.-R. Hsu, S.-H. Chen, C.-Y. Lin, AI4AMP: an antimicrobial peptide predictor using physicochemical property-based encoding method and deep learning. mSystems 6 (2021).

59. H. Lee, S. Lee, I. Lee, H. Nam, AMP-BERT: Prediction of antimicrobial peptide function based on a BERT model. Protein Sci. 32, e4529 (2023).

60. J. Yan, P. Bhadra, A. Li, P. Sethiya, L. Qin, H. K. Tai, K. H. Wong, S. W. I. Siu, Deep-AmPEP30: Improve short antimicrobial peptides prediction with deep learning. Mol. Ther. - Nucleic Acids 20, 882–894 (2020).

61. A. Bateman, M. J. Martin, S. Orchard, M. Magrane, R. Agivetova, S. Ahmad, E. Alpi, E. H. Bowler-Barnett, R. Britto, B. Bursteinas, H. Bye-A-Jee, R. Coetzee, A. Cukura, A. Da Silva, P. Denny, T. Dogan, T. G. Ebenezer, J. Fan, L. G. Castro, P. Garmiri, G. Georghiou, L. Gonzales, E. Hatton-Ellis, A. Hussein, A. Ignatchenko, G. Insana, R. Ishtiaq, P. Jokinen, V. Joshi, D. Jyothi, A. Lock, R. Lopez, A. Luciani, J. Luo, Y. Lussi, A. MacDougall, F. Madeira, M. Mahmoudy, M. Menchi, A. Mishra, K. Moulang, A. Nightingale, C. S. Oliveira, S. Pundir, G. Qi, S. Raj, D. Rice, M. R. Lopez, R. Saidi, J. Sampson, T. Sawford, E. Speretta, E. Turner, N. Tyagi, P. Vasudev, V. Volynkin, K. Warner, X. Watkins, R. Zaru, H. Zellner, A. Bridge, S. Poux, N. Redaschi, L. Aimo, G. Argoud-Puy, A. Auchincloss, K. Axelsen, P. Bansal, D. Baratin, M. C. Blatter, J. Bolleman, E. Boutet, L. Breuza, C. Casals-Casas, E. de Castro, K. C. Echioukh, E. Coudert, B. Cuche, M. Doche, D. Dornevil, A. Estreicher, M. L. Famiglietti, M. Feuermann, E. Gasteiger, S. Gehant, V. Gerritsen, A. Gos, N. Gruaz-Gumowski, U. Hinz, C. Hulo, N. Hyka-Nouspikel, F. Jungo, G. Keller, A. Kerhornou, V. Lara, P. Le Mercier, D. Lieberherr, T. Lombardot, X. Martin, P. Masson, A. Morgat, T. B. Neto, S. Paesano, I. Pedruzzi, S. Pilbout, L. Pourcel, M. Pozzato, M. Pruess, C. Rivoire, C. Sigrist, K. Sonesson, A. Stutz, S. Sundaram, M. Tognolli, L. Verbregue, C. H. Wu, C. N. Arighi, L. Arminski, C. Chen, Y. Chen, J. S. Garavelli, H. Huang, K. Laiho, P. McGarvey, D. A. Natale, K. Ross, C. R. Vinayaka, Q. Wang, Y. Wang, L. S. Yeh, J. Zhang, UniProt: the universal protein knowledgebase in 2021. Nucleic Acids Res. 49, D480–D489 (2021).

62. D. Osorio, P. Rondón-Villarreal, R. Torres, Peptides: A package for data mining of antimicrobial peptides. R J. 7, 4–14 (2015).

63. J. Jumper, R. Evans, A. Pritzel, T. Green, M. Figurnov, O. Ronneberger, K. Tunyasuvunakool, R. Bates, A. Žídek, A. Potapenko, A. Bridgland, C. Meyer, S. A. A. Kohl, A. J. Ballard, A. Cowie, B. Romera-Paredes, S. Nikolov, R. Jain, J. Adler, T. Back, S. Petersen, D. Reiman, E. Clancy, M. Zielinski, M. Steinegger, M. Pacholska, T. Berghammer, S. Bodenstein, D. Silver, O. Vinyals, A. W. Senior, K. Kavukcuoglu, P. Kohli, D. Hassabis, Highly accurate protein structure prediction with AlphaFold. Nature 596, 583–589 (2021).

64. P. J. A. Cock, T. Antao, J. T. Chang, B. A. Chapman, C. J. Cox, A. Dalke, I. Friedberg, T. Hamelryck, F. Kauff, B. Wilczynski, M. J. L. de Hoon, Biopython: freely available Python tools for computational molecular biology and bioinformatics. Bioinformatics 25, 1422–1423 (2009).

65. N. Mih, E. Brunk, K. Chen, E. Catoiu, A. Sastry, E. Kavvas, J. M. Monk, Z. Zhang, B. O. Palsson, ssbio: a Python framework for structural systems biology. Bioinformatics 34, 2155–2157 (2018).

66. H. Chen, F. Gu, Z. Huang, Improved Chou-Fasman method for protein secondary structure prediction. BMC Bioinformatics 7, 1–11 (2006).

67. R. Nagarajan, A. Archana, A. M. Thangakani, S. Jemimah, D. Velmurugan, M. M. Gromiha, PDBparam: online resource for computing structural parameters of proteins. Bioinform. Biol. Insights 10, 73–80 (2016).

68. W. Kabsch, C. Sander, Dictionary of protein secondary structure: pattern recognition of hydrogen-bonded and geometrical features. Biopolym. Orig. Res. Biomol. 22, 2577–2637 (1983).

69. W. G. Touw, C. Baakman, J. Black, T. A. H. Te Beek, E. Krieger, R. P. Joosten, G. Vriend, A series of PDB-related databanks for everyday needs. Nucleic Acids Res. 43, D364--D368 (2015).

70. V. Le Guilloux, P. Schmidtke, P. Tuffery, Fpocket: an open source platform for ligand pocket detection. BMC Bioinformatics 10, 1–11 (2009).

71. Y. Liang, S. Yang, L. Zheng, H. Wang, J. Zhou, S. Huang, L. Yang, Y. Zuo, Research progress of reduced amino acid alphabets in protein analysis and prediction. Comput. Struct. Biotechnol. J. (2022).

72. J. L. Figueroa, A. Redinbo, A. Panyala, S. Colby, M. L. Friesen, L. Tiemann, R. A. White, MerCat2: a versatile k-mer counter and diversity estimator for database-independent property analysis obtained from omics data. Bioinforma. Adv. 4 (2024).

73. F. Pedregosa, G. Varoquaux, A. Gramfort, M. Vincent, B. Thirion, O. Grisel, M. Blondel, P. Prettenhofer, R. Weiss, V. Dubourg, J. Vanderplas, A. Passos, D. Cournapeau, M. Brucher, M. Perrot, É. Duchesnay, Scikit-learn: machine learning in Python. J. Mach. Learn. Res. 12, 2825–2830 (2011).

74. Z. Lin, H. Akin, R. Rao, B. Hie, Z. Zhu, W. Lu, N. Smetanin, R. Verkuil, O. Kabeli, Y. Shmueli, others, Evolutionary-scale prediction of atomic-level protein structure with a language model. Science (80-. ). 379, 1123–1130 (2023).

75. R. de Jonge, H. P. Van Esse, K. Maruthachalam, M. D. Bolton, P. Santhanam, M. K. Saber, Z. Zhang, T. Usami, B. Lievens, K. V. Subbarao, B. P. H. J. Thomma, Tomato immune receptor Ve1 recognizes effector of multiple fungal pathogens uncovered by genome and RNA sequencing. Proc. Natl. Acad. Sci. U. S. A. 109, 5110–5115 (2012).

76. P. J. Kersey, J. E. Allen, I. Armean, S. Boddu, B. J. Bolt, D. Carvalho-Silva, M. Christensen, P. Davis, L. J. Falin, C. Grabmueller, others, Ensembl Genomes 2016: more genomes, more complexity. Nucleic Acids Res. 44, D574--D580 (2016).

77. H. Muraguchi, K. Umezawa, M. Niikura, M. Yoshida, T. Kozaki, K. Ishii, K. Sakai, M. Shimizu, K. Nakahori, Y. Sakamoto, C. Choi, C. Y. Ngan, E. Lindquist, A. Lipzen, A. Tritt, S. Haridas, K. Barry, I. V Grigoriev, P. J. Pukkila, Strand-specific RNA-Seq analyses of fruiting body development in *Coprinopsis cinerea*. PLoS One 10, e0141586 (2015).

78. I. V. Grigoriev, R. Nikitin, S. Haridas, A. Kuo, R. Ohm, R. Otillar, R. Riley, A. Salamov, X. Zhao, F. Korzeniewski, T. Smirnova, H. Nordberg, I. Dubchak, I. Shabalov, MycoCosm portal: Gearing up for 1000 fungal genomes. Nucleic Acids Res. 42, D699–D704 (2014).

79. G. Yildirir, J. Sperschneider, M. Malar C, E. C. H. Chen, W. Iwasaki, C. Cornell, N. Corradi, Long reads and Hi-C sequencing illuminate the two-compartment genome of the model arbuscular mycorrhizal symbiont *Rhizophagus irregularis*. New Phytol. 233, 1097–1107 (2022).

80. C. P. Cantalapiedra, A. Hern̗andez-Plaza, I. Letunic, P. Bork, J. Huerta-Cepas, eggNOG-mapper v2: functional annotation, orthology assignments, and domain prediction at the metagenomic scale. Mol. Biol. Evol. 38, 5825–5829 (2021).

81. J. Huerta-Cepas, D. Szklarczyk, D. Heller, A. Hernández-Plaza, S. K. Forslund, H. Cook, D. R. Mende, I. Letunic, T. Rattei, L. J. Jensen, C. Von Mering, P. Bork, eggNOG 5.0: a hierarchical, functionally and phylogenetically annotated orthology resource based on 5090 organisms and 2502 viruses. Nucleic Acids Res. 47, D309–D314 (2019).

82. J. Zheng, Q. Ge, Y. Yan, X. Zhang, L. Huang, Y. Yin, dbCAN3: automated carbohydrate-active enzyme and substrate annotation. Nucleic Acids Res., gkad328 (2023).

83. M. Bernhofer, B. Rost, TMbed: transmembrane proteins predicted through language model embeddings. BMC Bioinformatics 23, 1–19 (2022).

84. D. M. Emms, S. Kelly, OrthoFinder: Phylogenetic orthology inference for comparative genomics. Genome Biol. 20, 1–14 (2019).

85. F. Mesny, S. Miyauchi, T. Thiergart, B. Pickel, L. Atanasova, M. Karlsson, B. Hüttel, K. W. Barry, S. Haridas, C. Chen, D. Bauer, W. Andreopoulos, J. Pangilinan, K. LaButti, R. Riley, A. Lipzen, A. Clum, E. Drula, B. Henrissat, A. Kohler, I. V. Grigoriev, F. M. Martin, S. Hacquard, Genetic determinants of endophytism in the *Arabidopsis* root mycobiome. Nat. Commun. 12, 1–15 (2021).

86. D. M. Emms, S. Kelly, STAG: Species tree inference from all genes. bioRxiv, 267914 (2018).

87. F. Mesny, phylorep v0.1. (2023). 10.5281/ZENODO.10142123.

88. M. N. Price, P. S. Dehal, A. P. Arkin, FastTree 2 – Approximately maximum-likelihood trees for large alignments. PLoS One 5, e9490 (2010).

89. G. L. Fiorin, A. Sanchéz-Vallet, D. P. de T. Thomazella, P. F. V. do Prado, L. C. do Nascimento, A. V. de O. Figueira, B. P. H. J. Thomma, G. A. G. Pereira, P. J. P. L. Teixeira, Suppression of plant immunity by fungal chitinase-like effectors. Curr. Biol. 28, 3023–3030.e5 (2018).

90. H. Tian, C. I. MacKenzie, L. Rodriguez-Moreno, G. C. M. van den Berg, H. Chen, J. J. Rudd, J. R. Mesters, B. P. H. J. Thomma, Three LysM effectors of *Zymoseptoria tritici* collectively disarm chitin-triggered plant immunity. Mol. Plant Pathol. 22, 683–693 (2021).

91. C. A. Schneider, W. S. Rasband, K. W. Eliceiri, NIH Image to ImageJ: 25 years of image analysis. Nat. Methods 9, 671–675 (2012).

92. K. Katoh, D. M. Standley, MAFFT multiple sequence alignment software version 7: Improvements in performance and usability. Mol. Biol. Evol. 30, 772–780 (2013).

93. P. Jones, D. Binns, H. Y. Chang, M. Fraser, W. Li, C. McAnulla, H. McWilliam, J. Maslen, A. Mitchell, G. Nuka, S. Pesseat, A. F. Quinn, A. Sangrador-Vegas, M. Scheremetjew, S. Y. Yong, R. Lopez, S. Hunter, InterProScan 5: genome-scale protein function classification. Bioinformatics 30, 1236–1240 (2014).

94. I. Letunic, P. Bork, Interactive Tree of Life (iTOL) v6: recent updates to the phylogenetic tree display and annotation tool. Nucleic Acids Res. 52, W78–W82 (2024).

95. O. Emanuelsson, H. Nielsen, G. Von Heijne, ChloroP, a neural network-based method for predicting chloroplast transit peptides and their cleavage sites. Protein Sci. 8, 978–984 (1999).

96. W. Punt, A. Kraege, …, B. P. H. J. Thomma, An antimicrobial effector from Verticillium dahliae differentially contributes to virulence and differentially impacts tomato microbiota across natural soils. In Prep.

97. J. Li, L. Faino, G. L. Fiorin, S. Bashyal, A. Schaveling, C. van den Berg, M. F. Seidl, B. PHJ Thomma, B. Thomma, A single Verticillium dahliae effector determines pathogenicity on tomato by targeting auxin response factors. bioRxiv, 2022.11.22.517554 (2022).

98. L. Rehman, X. Su, H. Guo, X. Qi, H. Cheng, Protoplast transformation as a potential platform for exploring gene function in *Verticillium dahliae*. BMC Biotechnol. 16, 1–9 (2016).

99. T. Leisen, F. Bietz, J. Werner, A. Wegner, U. Schaffrath, D. Scheuring, F. Willmund, A. Mosbach, G. Scalliet, M. Hahn, CRISPR/Cas with ribonucleoprotein complexes and transiently selected telomere vectors allows highly efficient marker-free and multiple genome editing in *Botrytis cinerea*. PLOS Pathog. 16, e1008326 (2020).

100. X. Guo, X. Zhang, Y. Qin, Y. X. Liu, J. Zhang, N. Zhang, K. Wu, B. Qu, Z. He, X. Wang, X. Zhang, S. Hacquard, X. Fu, Y. Bai, Host-associated quantitative abundance profiling reveals the microbial load variation of root microbiome. Plant Commun. 1 (2020).

101. J. M. Kremer, B. C. Paasch, D. Rhodes, C. Thireault, J. E. Froehlich, P. Schulze-Lefert, J. M. Tiedje, S. Y. He, FlowPot axenic plant growth system for microbiota research. bioRxiv, 254953 (2018).

102. B. Schlesier, F. Bréton, H. P. Mock, A hydroponic culture system for growing *Arabidopsis thaliana* plantlets under sterile conditions. Plant Mol. Biol. Report. 21, 449–456 (2003).

103. P. Virtanen, R. Gommers, T. E. Oliphant, M. Haberland, T. Reddy, D. Cournapeau, E. Burovski, P. Peterson, W. Weckesser, J. Bright, S. J. van der Walt, M. Brett, J. Wilson, K. J. Millman, N. Mayorov, A. R. J. Nelson, E. Jones, R. Kern, E. Larson, C. J. Carey, İ. Polat, Y. Feng, E. W. Moore, J. VanderPlas, D. Laxalde, J. Perktold, R. Cimrman, I. Henriksen, E. A. Quintero, C. R. Harris, A. M. Archibald, A. H. Ribeiro, F. Pedregosa, P. van Mulbregt, A. Vijaykumar, A. Pietro Bardelli, A. Rothberg, A. Hilboll, A. Kloeckner, A. Scopatz, A. Lee, A. Rokem, C. N. Woods, C. Fulton, C. Masson, C. Häggström, C. Fitzgerald, D. A. Nicholson, D. R. Hagen, D. V. Pasechnik, E. Olivetti, E. Martin, E. Wieser, F. Silva, F. Lenders, F. Wilhelm, G. Young, G. A. Price, G. L. Ingold, G. E. Allen, G. R. Lee, H. Audren, I. Probst, J. P. Dietrich, J. Silterra, J. T. Webber, J. Slavič, J. Nothman, J. Buchner, J. Kulick, J. L. Schönberger, J. V. de Miranda Cardoso, J. Reimer, J. Harrington, J. L. C. Rodríguez, J. Nunez-Iglesias, J. Kuczynski, K. Tritz, M. Thoma, M. Newville, M. Kümmerer, M. Bolingbroke, M. Tartre, M. Pak, N. J. Smith, N. Nowaczyk, N. Shebanov, O. Pavlyk, P. A. Brodtkorb, P. Lee, R. T. McGibbon, R. Feldbauer, S. Lewis, S. Tygier, S. Sievert, S. Vigna, S. Peterson, S. More, T. Pudlik, T. Oshima, T. J. Pingel, T. P. Robitaille, T. Spura, T. R. Jones, T. Cera, T. Leslie, T. Zito, T. Krauss, U. Upadhyay, Y. O. Halchenko, Y. Vázquez-Baeza, SciPy 1.0: fundamental algorithms for scientific computing in Python. Nat. Methods 2020 173 17, 261–272 (2020).

104. S. Seabold, J. Perktold, “statsmodels: Econometric and statistical modeling with python” in 9th {Python} in {Science} {Conference} (2010).

105. S. Graves, H.-P. Piepho, L. Selzer, multcompView: Visualizations of paired comparisons. (2024). https://github.com/lselzer/multcompview.

106. G. Wang, X. Li, Z. Wang, APD3: the antimicrobial peptide database as a tool for research and education. Nucleic Acids Res. 44, D1087–D1093 (2016).

107. L. V. Hedges, Distribution theory for Glass’s estimator of effect size and related estimators. J. Educ. Stat. 6, 107–128 (1981).

