## Supplementary Material for "Plant-associated fungi co-opt ancient antimicrobials for host manipulation"

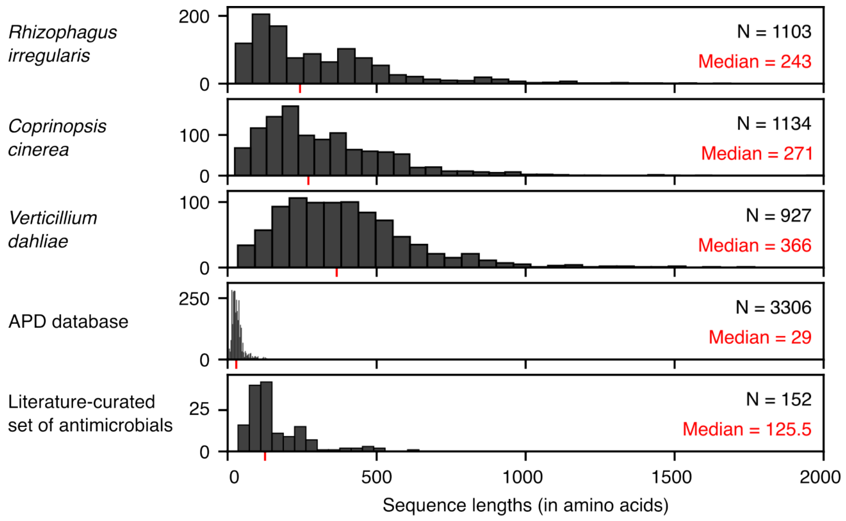

**Fig. S1. Protein sequence lengths in three fungal secretomes, the AMP database APD and a literature-curated set of antimicrobials.** The top three histograms show mature sequence lengths (in number of amino acids) of secreted proteins predicted with SignalP (*55*) in three fungi selected based on their distance in the tree of life and their distinct lifestyles: the Glomeromycota mycorrhizal fungus *Rhizophagus irregularis*, the Basidiomycete saprophyte *Coprinopsis cinerea*, and the Ascomycete plant pathogen *Verticillium dahliae*. The fourth histogram shows the length of AMPs in the APD database (*106*), which was previously used to train published antimicrobial peptide predictors. The bottom histogram shows sequence lengths in our newly curated set of antimicrobial proteins, used as a positive training set to develop AMAPEC.

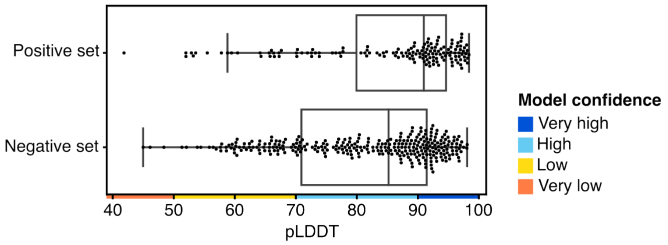

**Fig. S2. Confidence of predicted structures in the training datasets.** Boxplots showing the distribution of mean pLDDT confidence scores of AlphaFold-predicted protein structures in the positive and negative training sets. The color code depicting model confidence originates from the AlphaFold documentation (*63*).

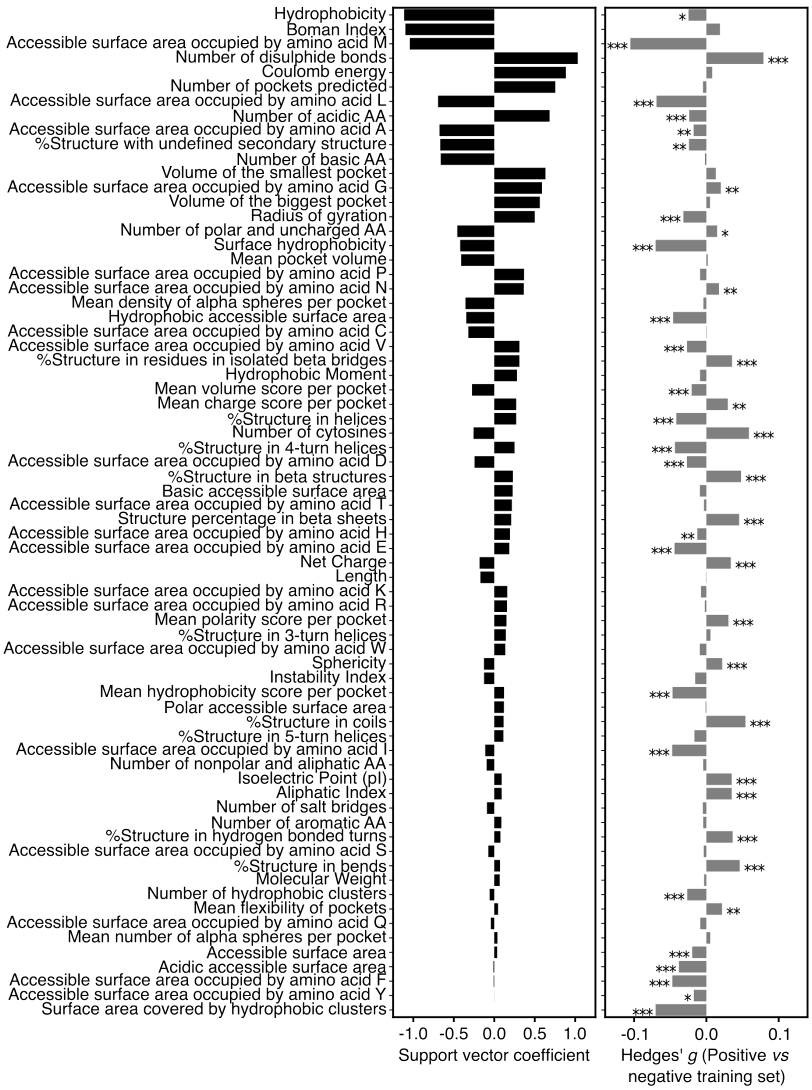

**Fig. S3. Physicochemical properties of proteins and their importance for antimicrobial activity prediction.** Physicochemical properties implemented in the AMAPEC training pipeline (Fig. 1, C) are listed and ranked according to their importance for our Support Vector Machines (SVM) classifier (vector weights). The left barplot (black) shows support vector coefficients, representing vector weights and orientation. The right barplot (grey) shows the results of an enrichment analysis testing for significant differences between values in the positive and in the negative training set. This analysis was conducted by Mann-Whitney U test and Benjamini-Hochberg correction (FDR values depicted with asterisks: *: ≤0.05; **≤0.01; ***≤0.001) and we additionally calculated standard effect sizes (Hedges' g, (*107*)).

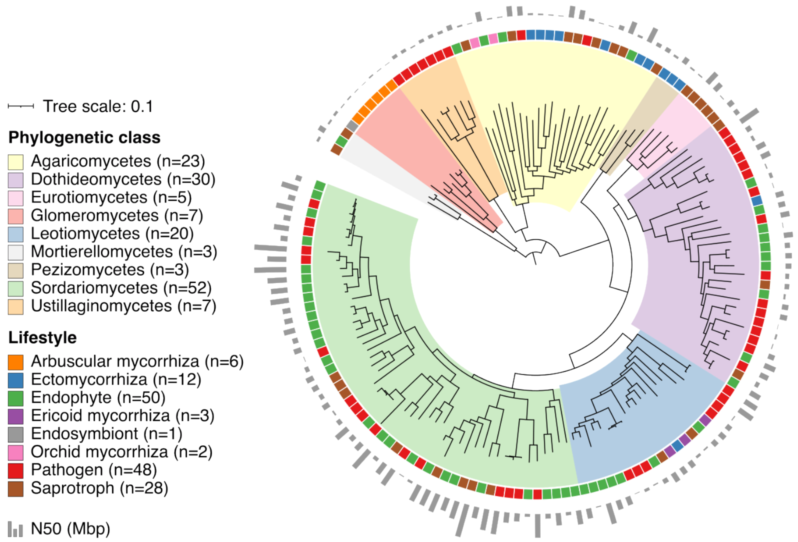

**Fig. S4. Description of the dataset of 150 fungal genomes used for comparative genomics.** Phylogenomic tree calculated on total sets of proteins from the selected 150 fungi (STAG method (*86*) implemented in OrthoFinder (*84*)). Color ranges on the phylogenomic tree highlight phylogenetic classes and fungal lifestyles are indicated. A barplot shows the N50 values, reflecting overall genome quality and fragmentation.

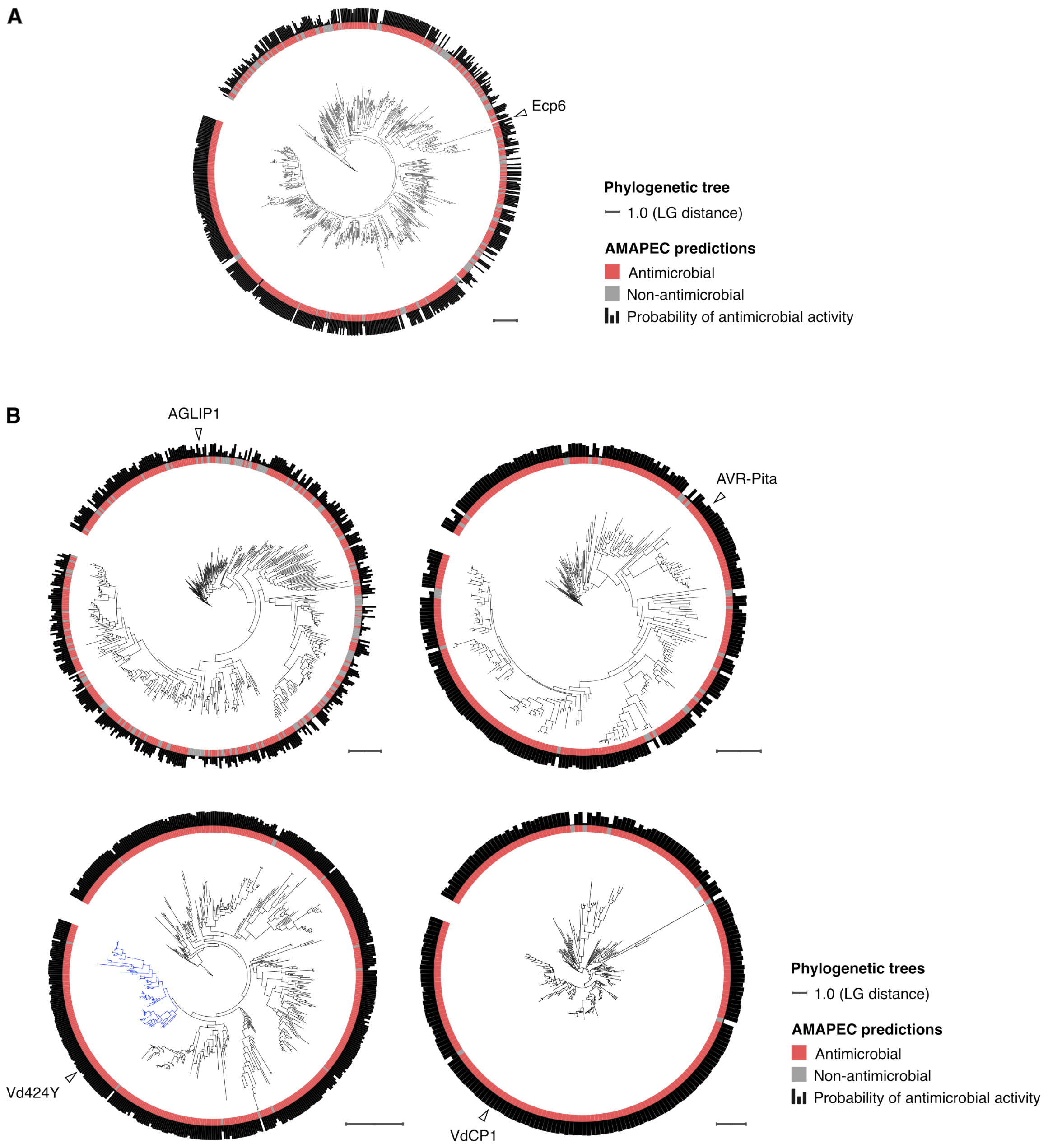

**Fig. S5. Phylogenies of five secreted protein families, including reference effectors, with antimicrobial activity prediction for each family member.** (**A**) Phylogeny and antimicrobial activity prediction calculated on LysM effectors annotated in the fungal dataset of 150 genomes and additionally in *Cladosporium fulvum*, which encodes the reference Ecp6 effector (*37*). (**B**) Phylogenies of the families of AGLIP1, AVR-Pita, Vd424Y and VdCP1 as defined by orthology prediction in the dataset of 150 fungal genomes. All five phylogenies were calculated by sequence alignment of mature sequence proteins with MAFFT (*92*), and phylogenetic reconstruction with IQ-TREE (*53*) with maximum-likelihood model 'LG'. In the family tree of Vd424Y, the clade depicted in blue was considered as the Vd424Y subfamily and was further analyzed in Fig. 4, A and fig. S16.

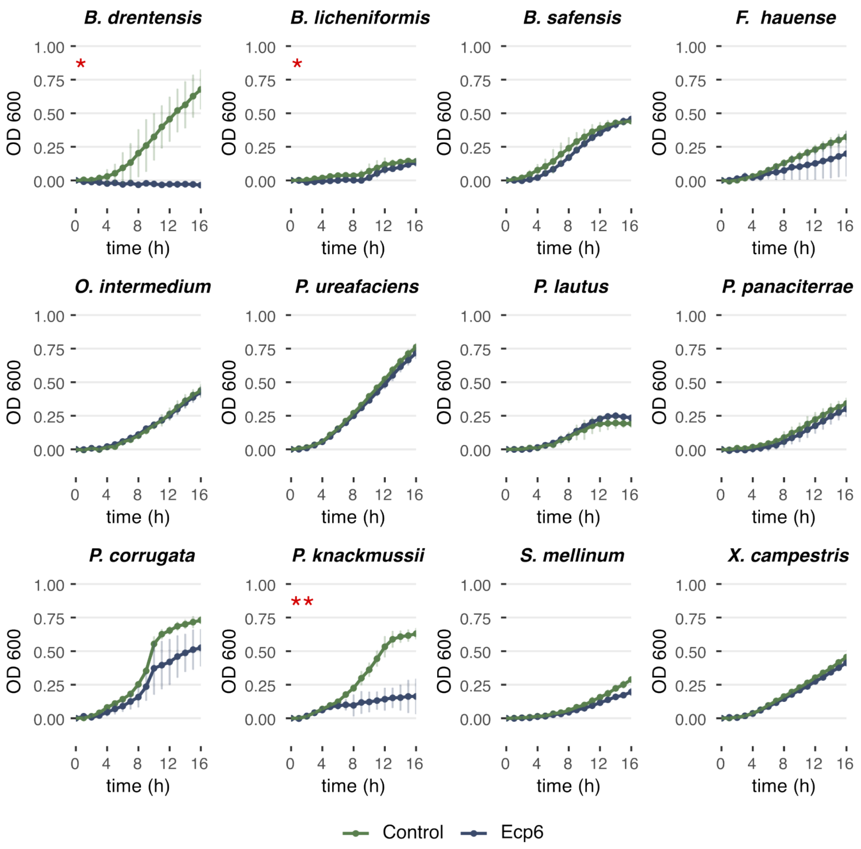

**Fig. S6. Selective antibacterial activity displayed *in vitro* by the Ecp6 effector protein of the tomato leaf mold pathogen *Cladosporium fulvum*.** Absorbance measurements (at wavelength 600 nm) over 16 hours of bacterial cultivation in presence and absence of 8 µM of heterologously produced effector protein. Each growth curve represents to the mean $OD_{600}$ over 3 independent replicates and error bars correspond to the standard deviation. The assay was performed on a phylogenetically diverse set of 12 bacterial isolates, which species-level phylogeny can be seen on Fig. 3, C. Asterisks highlight significance (p-value < 0.05) of a Student's T-tests computed on area-under-curve values comparing bacterial growth in presence and absence of effector protein: ***: $P < 0.001$, **: $P < 0.01$, *: $P < 0.05$.

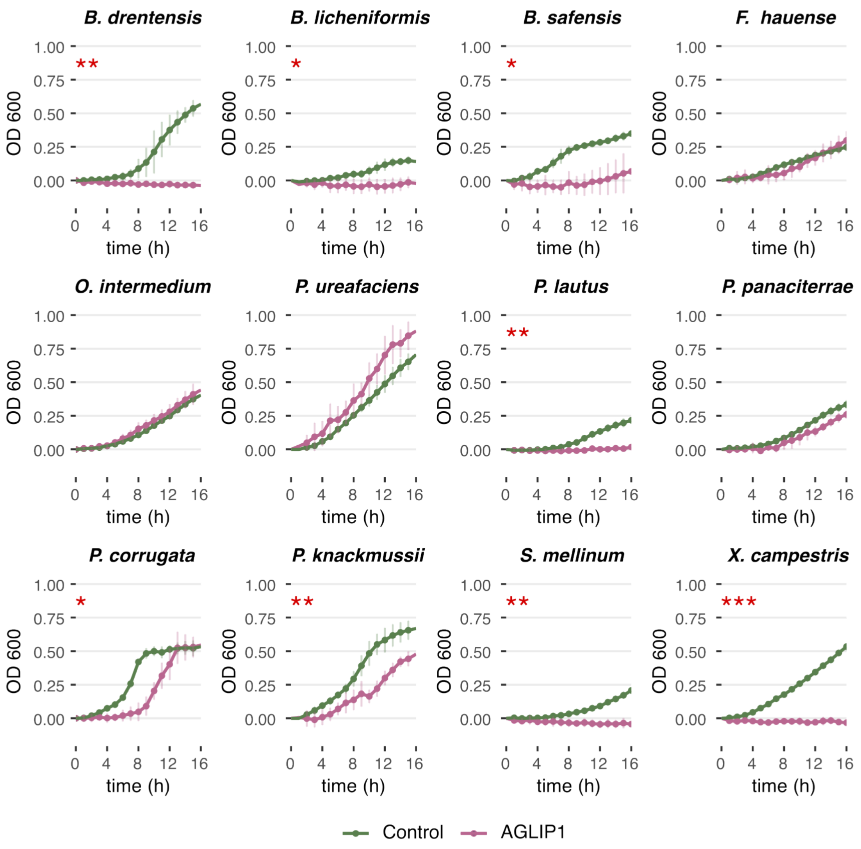

**Fig. S7. Selective antibacterial activity displayed *in vitro* displayed by the AGLIP1 effector protein of the root rot pathogen *Rhizoctonia solani*.** Absorbance measurements (at wavelength 600 nm) over 16 hours of bacterial cultivation in presence and absence of 8 µM of heterologously produced effector protein. Each growth curve represents to the mean $OD_{600}$ over 3 independent replicates and error bars correspond to the standard deviation. The assay was performed on a phylogenetically diverse set of 12 bacterial isolates, which species-level phylogeny can be seen on Fig. 3, C. Asterisks highlight significance (p-value < 0.05) of a Student's T-tests computed on area-under-curve values comparing bacterial growth in presence and absence of effector protein: ***: $P < 0.001$, **: $P < 0.01$, *: $P < 0.05$.

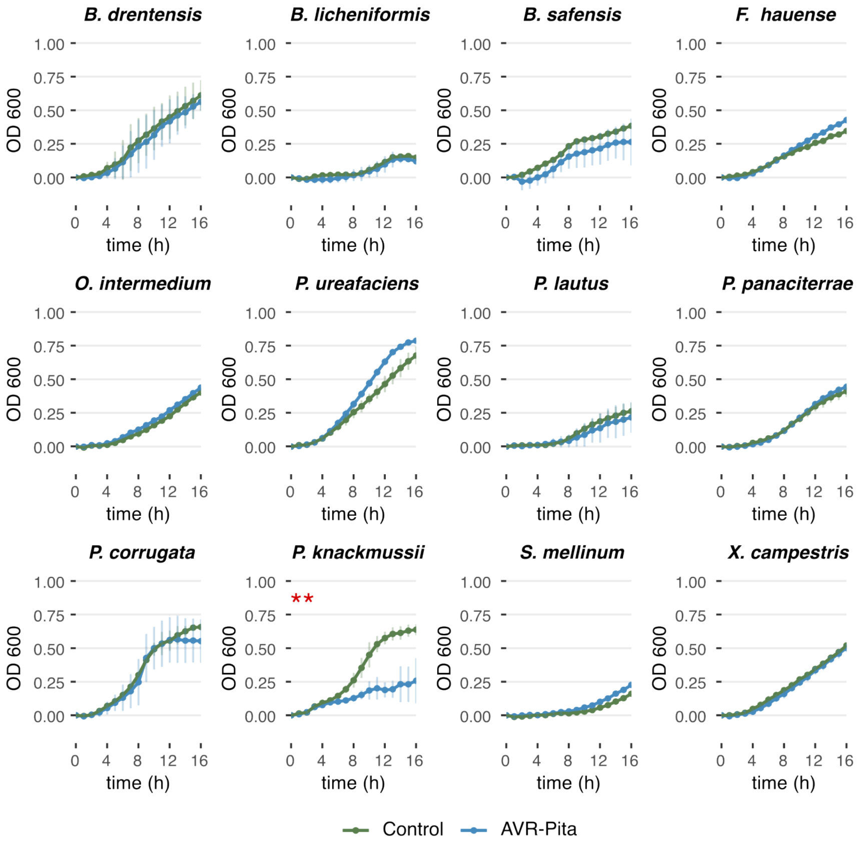

**Fig. S8. Selective antibacterial activity displayed *in vitro* by the AVR-Pita effector protein of the rice blast pathogen *Magnaporthe oryzae*.** Absorbance measurements (at wavelength 600 nm) over 16 hours of bacterial cultivation in presence and absence of 8 µM of heterologously produced effector protein. Each growth curve represents to the mean $OD_{600}$ over 3 independent replicates and error bars correspond to the standard deviation. The assay was performed on a phylogenetically diverse set of 12 bacterial isolates, which species-level phylogeny can be seen on Fig. 3, C. Asterisks highlight significance (p-value < 0.05) of a Student's T-tests computed on area-under-curve values comparing bacterial growth in presence and absence of effector protein: ***: $P < 0.001$, **: $P < 0.01$, *: $P < 0.05$.

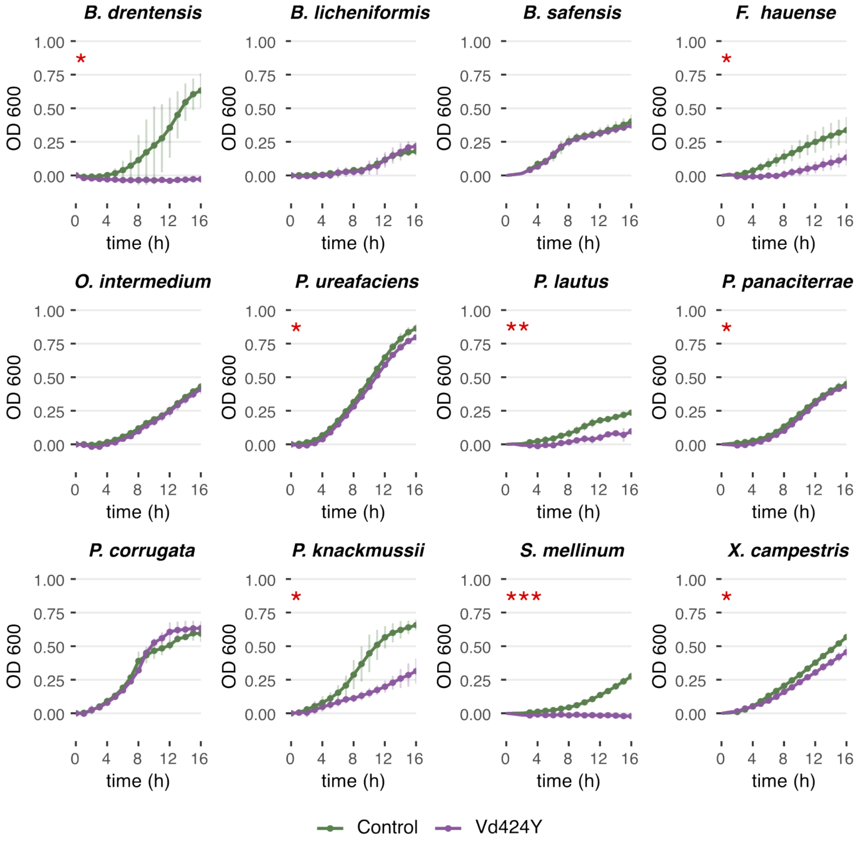

**Fig. S9. Selective antibacterial activity displayed *in vitro* by the Vd424Y effector protein of the vascular wilt pathogen *Verticillium dahliae*.** Absorbance measurements (at wavelength 600 nm) over 16 hours of bacterial cultivation in presence and absence of 8 µM of heterologously produced effector protein. Each growth curve represents to the mean $OD_{600}$ over 3 independent replicates and error bars correspond to the standard deviation. The assay was performed on a phylogenetically diverse set of 12 bacterial isolates, which species-level phylogeny can be seen on Fig. 3, C. Asterisks highlight significance (p-value < 0.05) of a Student's T-tests computed on area-under-curve values comparing bacterial growth in presence and absence of effector protein: ***: $P < 0.001$, **: $P < 0.01$, *: $P < 0.05$.

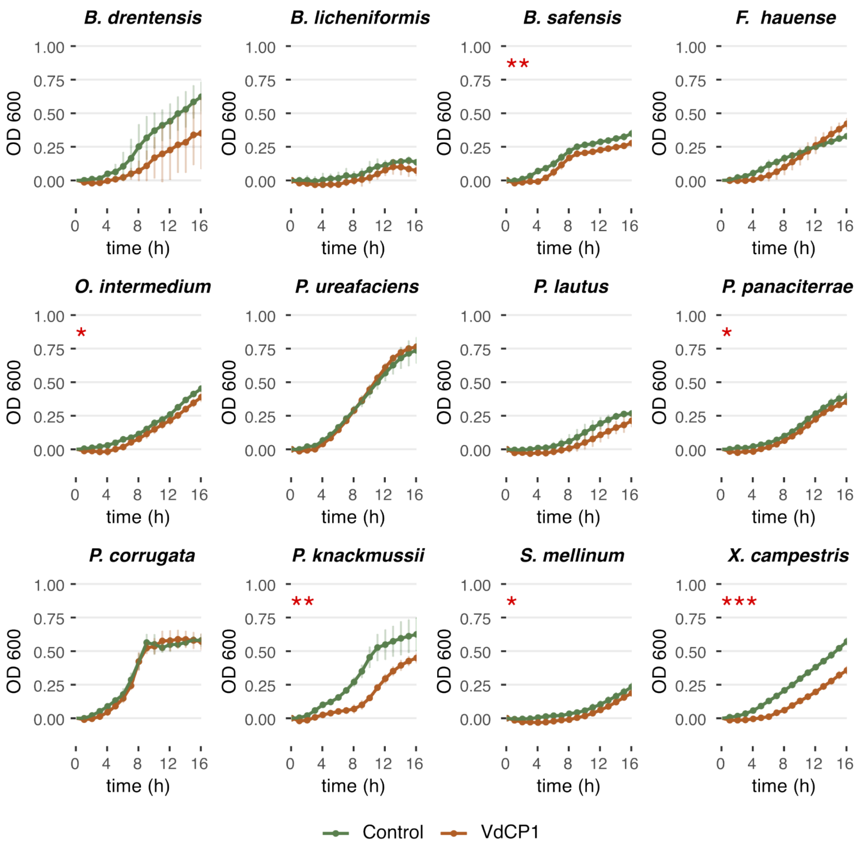

**Fig. S10. Selective antibacterial activity displayed *in vitro* by the VdCP1 effector protein of the vascular wilt pathogen *Verticillium dahliae*.** Absorbance measurements (at wavelength 600 nm) over 16 hours of bacterial cultivation in presence and absence of 8 µM of heterologously produced effector protein. Each growth curve represents to the mean $OD_{600}$ over 3 independent replicates and error bars correspond to the standard deviation. The assay was performed on a phylogenetically diverse set of 12 bacterial isolates, which species-level phylogeny can be seen on Fig. 3, C. Asterisks highlight significance (p-value < 0.05) of a Student's T-tests computed on area-under-curve values comparing bacterial growth in presence and absence of effector protein: ***: $P < 0.001$, **: $P < 0.01$, *: $P < 0.05$.

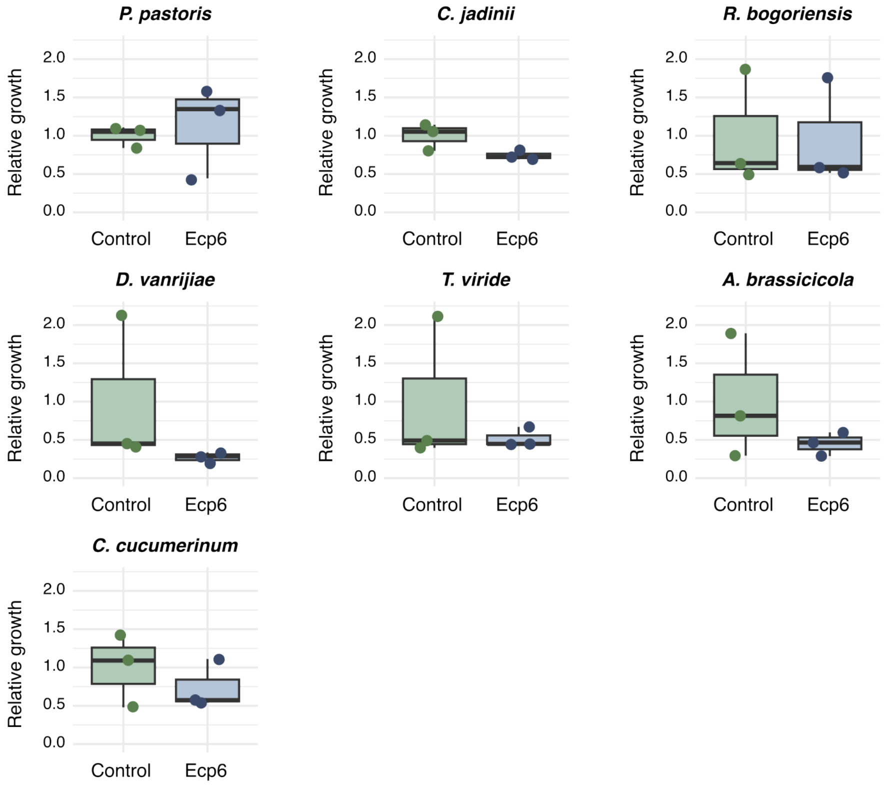

**Fig. S11. Absence of antifungal activity displayed *in vitro* by the Ecp6 effector protein of the tomato leaf mold pathogen *Cladosporium fulvum*.** Normalized fungal areas measured on microscopy photographs of growth medium, after 16 hours of fungal culture in presence and absence of 8 µM of heterologously produced effector protein. The assay was performed on a phylogenetically diverse set of seven fungal isolates, which species-level phylogeny can be seen on Fig. 3, D. Asterisks highlight significance (p-value < 0.05) of a Student's T-tests computed comparing fungal growth in presence and absence of effector protein: ***: $P < 0.001$, **: $P < 0.01$, *: $P < 0.05$.

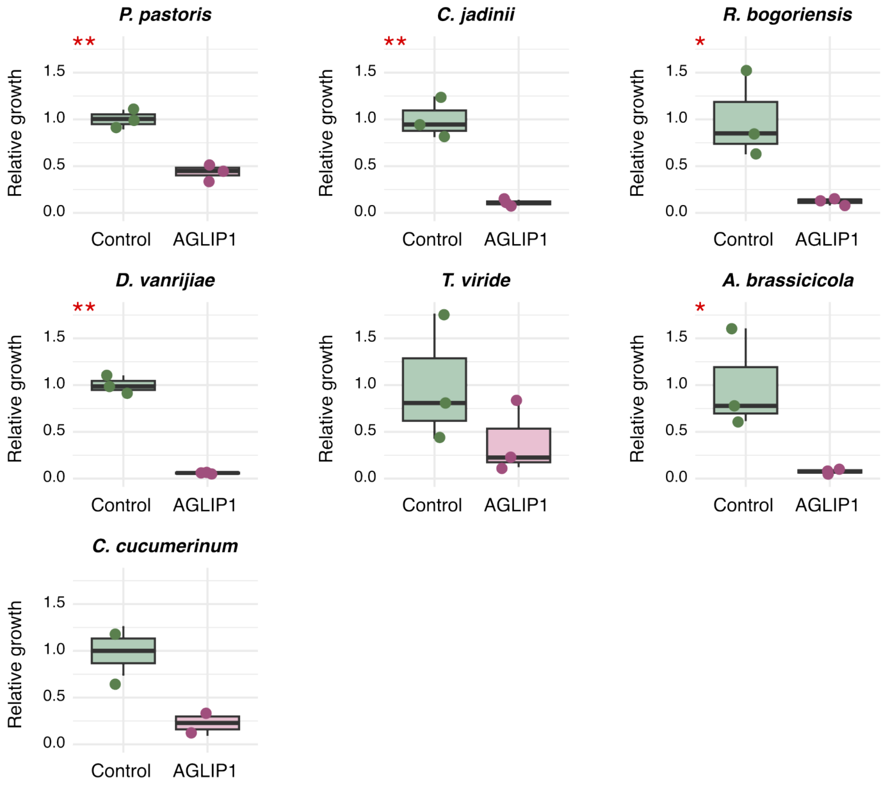

**Fig. S12. Antifungal activity displayed *in vitro* by the AGLIP1 effector protein of the root rot pathogen *Rhizoctonia solani*.** Normalized fungal areas measured on microscopy photographs of growth medium, after 16 hours of fungal culture in presence and absence of 8 µM of heterologously produced effector protein. The assay was performed on a phylogenetically diverse set of seven fungal isolates, which species-level phylogeny can be seen on Fig. 3, D. Asterisks highlight significance (p-value < 0.05) of a Student's T-tests computed comparing fungal growth in presence and absence of effector protein: ***: $P < 0.001$, **: $P < 0.01$, *: $P < 0.05$.

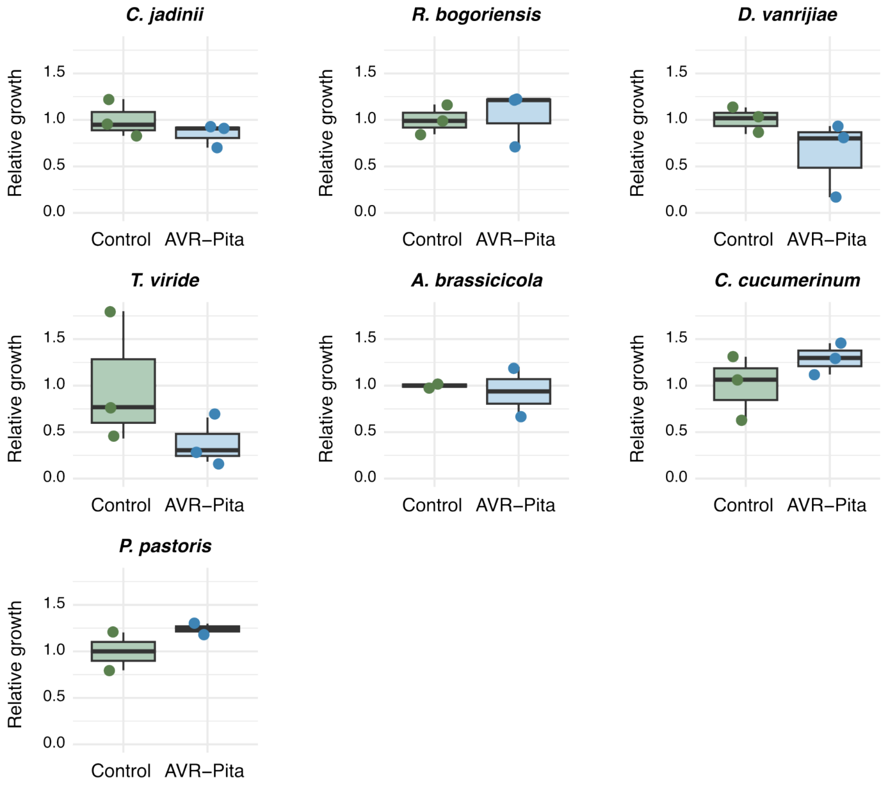

**Fig. S13. Absence of antifungal activity displayed *in vitro* by the AVR-Pita effector protein of the rice blast pathogen *Magnaporthe oryzae*.** Normalized fungal areas measured on microscopy photographs of growth medium, after 16 hours of fungal culture in presence and absence of 8 µM of heterologously produced effector protein. The assay was performed on a phylogenetically diverse set of seven fungal isolates, which species-level phylogeny can be seen on Fig. 3, D. Asterisks highlight significance (p-value < 0.05) of a Student's T-tests computed comparing fungal growth in presence and absence of effector protein: ***: $P < 0.001$, **: $P < 0.01$, *: $P < 0.05$.

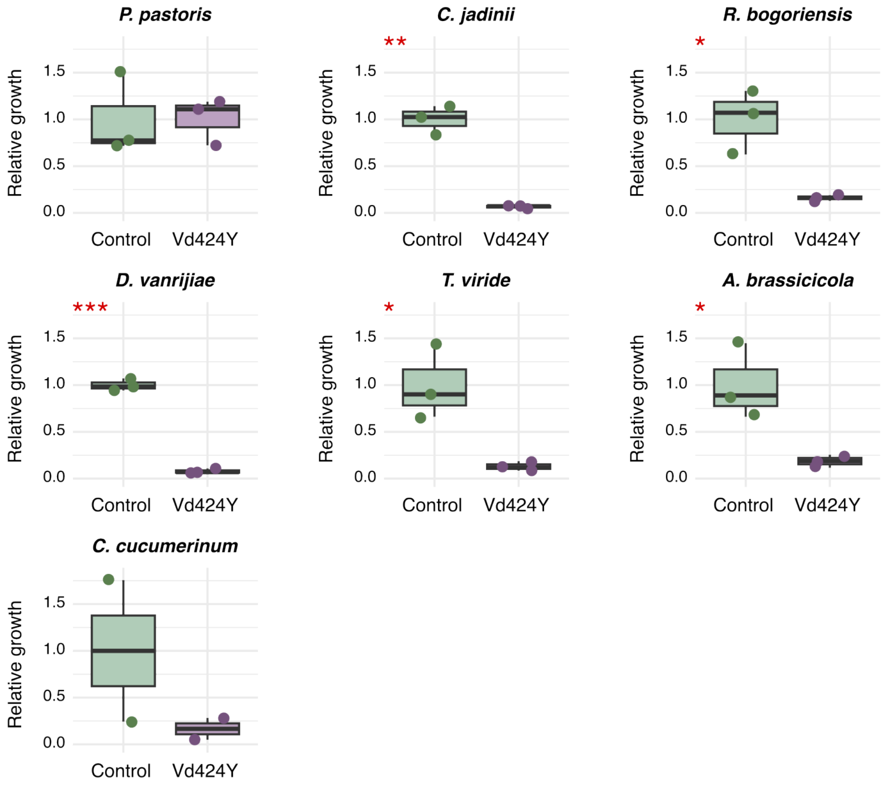

**Fig. S14. Antifungal activity displayed *in vitro* by the Vd424Y effector protein of the vascular wilt pathogen *Verticillium dahliae*.** Normalized fungal areas measured on microscopy photographs of growth medium, after 16 hours of fungal culture in presence and absence of 8 µM of heterologously produced effector protein. The assay was performed on a phylogenetically diverse set of seven fungal isolates, which species-level phylogeny can be seen on Fig. 3, D. Asterisks highlight significance (p-value < 0.05) of a Student's T-tests computed comparing fungal growth in presence and absence of effector protein: ***: $P < 0.001$, **: $P < 0.01$, *: $P < 0.05$.

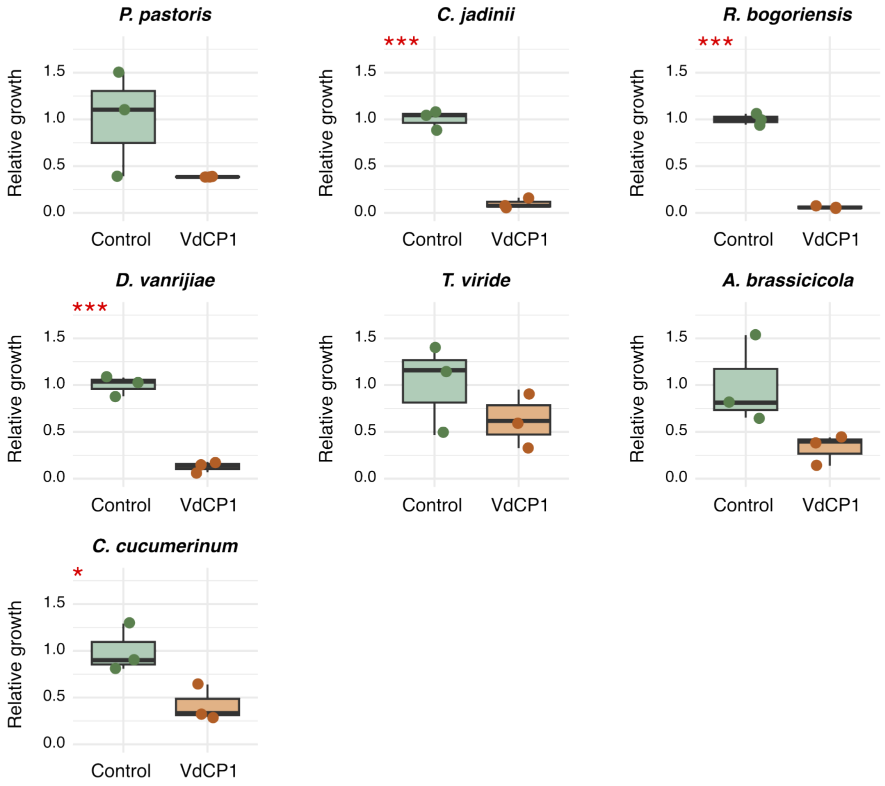

**Fig. S15. Antifungal activity displayed *in vitro* by the VdCP1 effector protein of the vascular wilt pathogen *Verticillium dahliae*.** Normalized fungal areas measured on microscopy photographs of growth medium, after 16 hours of fungal culture in presence and absence of 8 µM of heterologously produced effector protein. The assay was performed on a phylogenetically diverse set of seven fungal isolates, which species-level phylogeny can be seen on Fig. 3, D. Asterisks highlight significance (p-value < 0.05) of a Student's T-tests computed comparing fungal growth in presence and absence of effector protein: ***: $P < 0.001$, **: $P < 0.01$, *: $P < 0.05$.

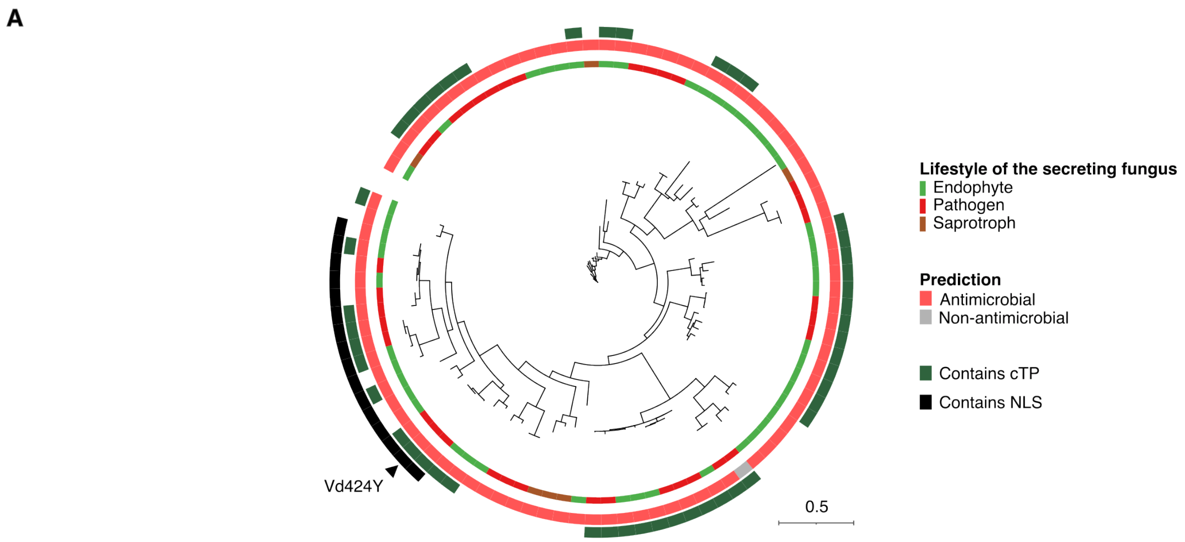

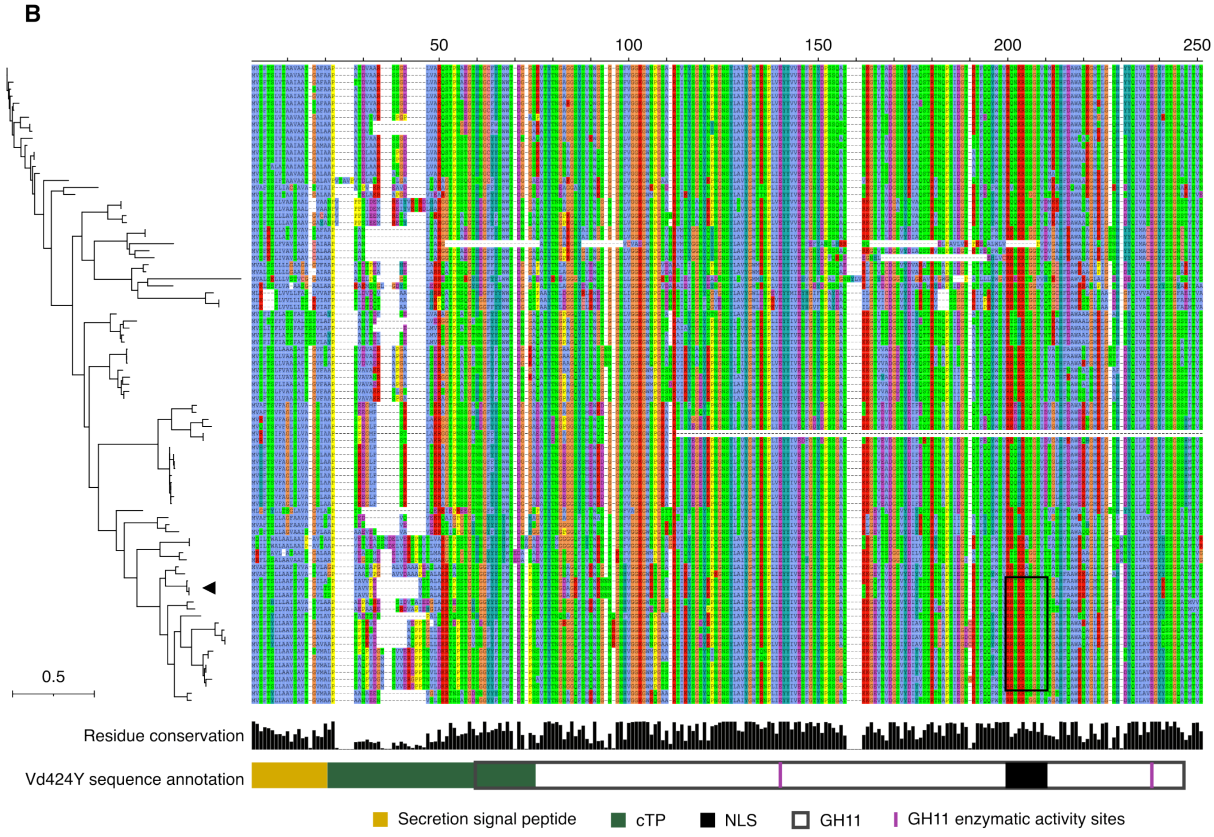

**Fig. S16. Evolution of Vd424Y sequence features.** (**A**) Maximum-likelihood phylogenetic tree (computed with IQ-TREE (*53*), model 'LG') of Vd424Y homologs in a dataset of 150 fungal genomes. The set of proteins represent those occurring in the same subfamily (large clade) as Vd424Y, as identified in the total family phylogenetic tree (fig. S5). The tree was manually rooted at the protein identified as an optimal outgroup by IQ-TREE and annotated with the lifestyles of the fungus secreting each protein, the results of antimicrobial activity as well as the occurrence of a chloroplast transit peptide (cTP) and nuclear localization signal (NLS) annotated using ChloroP (*95*) and cNLS Mapper (*54*), respectively. (**B**) The same phylogenetic tree as on panel A is displayed on the left, with a black triangle indicating the location of Vd424Y. On the right, the protein sequence alignment (generated with MAFFT (*92*)) presents sequence variation in the Vd424Y subfamily. A black rectangle circumscribes annotated NLS motifs in the sequences. At the bottom, a barplot presents the conservation of amino acids at each position in the protein sequence. Below, a diagram shows the organization in functional domains of the Vd424Y sequence, as characterized previously (*32*).

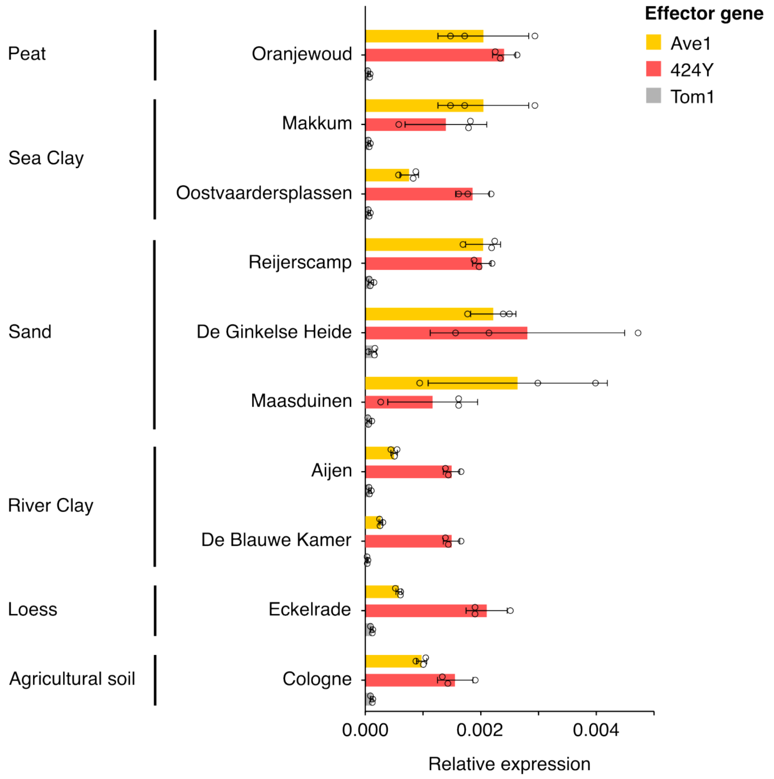

**Fig. S17. Expression of the *V. dahliae* 424Y-encoding gene in a diverse set of soils.** Real-time PCR measurements of effector gene expression in a diverse set of 10 soils (*96*), classified by soil type (left). In addition to the 424Y-encoding gene, effector genes Ave1 (*7*) and Tom1 (*97*) were studied and serve as positive and negative controls, respectively.

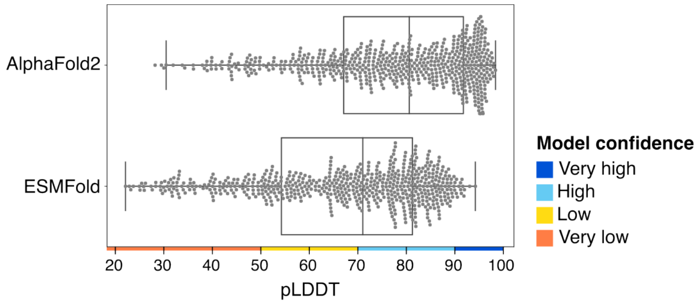

**Fig. S18. Confidence of AlphaFold2- and ESMFold-predicted structures for secreted proteins of *Verticillium dahliae*.** Boxplots showing the distribution of mean pLDDT confidence scores of AlphaFold2 (*63*)- and ESMFold (*74*)-predicted structures for 626 non-CAZyme secreted proteins of *Verticillium dahliae*. While the secretome of *V. dahliae* includes 635 non-CAZyme proteins, AlphaFold2 failed at predicting the structures of nine of these proteins, which were therefore excluding from this analysis. The color code depicting model confidence originates from the documentation of AlphaFold2.

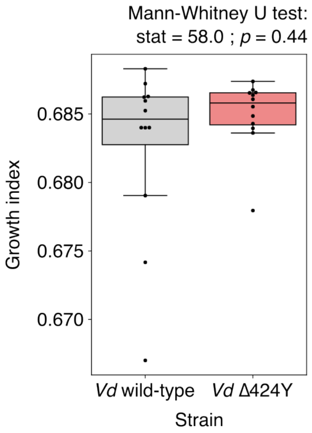

**Fig. S19. Growth of *Verticillium dahliae* wild-type and Δ424Y mutant *in vitro*.** Boxplot showing growth indices of *Verticillium dahliae* JR2 wild-type and Δ424Y calculated from quantitative PCR Ct values (see Material and methods for details) after 48 hours in growth medium. In total, 12 samples per condition corresponding to 12 independent fungal cultures over 3 biological replicates were analyzed. Results of a Mann-Whitney U test revealing no significant difference between the wild-type and the mutant strain are written on top of the figure.

### List and legends of supplementary tables

**Table S1. Description of the literature-curated set of antimicrobial proteins.**
For each protein in the set, the table provides (1) a reference identifier; (2) the name of the protein in literature; (3) the reported antimicrobial activity in literature; (4) the phylogenetic group of organisms by which it is encoded; (5) the species producing the protein; (6) the publication that described the antimicrobial activity of this protein; (7) whether a secretion signal was identified by SignalP (*55*) and removed from the protein sequence; (8) the pLDDT confidence score for the AlphaFold2 (*63*)-predicted structure.

**Table S2. Description of the negative training set of presumably non-antimicrobial proteins.**
For each protein in the set, the table provides (1) a reference identifier; (2) the functional description of the protein in the UniProt database (*61*); (3) the phylogenetic group of organisms encoding the protein; (4) the species producing the protein; (5) the UniProt entry identifier; (6) whether a secretion signal was identified by SignalP (*55*) and removed from the protein sequence; (7) the pLDDT confidence score for the AlphaFold2 (*63*)-predicted structure.

**Table S3. Properties and k-mers describing protein physicochemistry used to predict antimicrobial activity.**
List of 70 properties calculated from protein sequences and structures, with as a reference, the method implementing the calculation or publication introducing the formula. Additionally, the list includes 6 k-mers (in a reduced amino acid alphabet designed based on amino acid properties, see Material and Methods) found to be over- or under-represented in antimicrobial protein sequences. All 76 properties are used by the predictor to classify a protein as antimicrobial or non-antimicrobial.

**Table S4. Prediction of antimicrobial activity for six recently characterized fungal antimicrobial proteins.**
List of six recently characterized fungal antimicrobial proteins with their organism of origin, reference publication and sequence, together with the results of antimicrobial activity prediction after structure prediction with ESMFold (*74*).

**Table S5. Functional annotation of the secretome of the arbuscular mycorrhizal glomeromycete *Rhizophagus irregularis* and antimicrobial activity prediction results.**
Secretome functional annotation outputs of emapper (*80*), transmembrane protein prediction by TMBed (*83*) and carbohydrate-active enzyme annotation from dbCAN (*82*), together with the results of antimicrobial activity prediction with AMAPEC.

**Table S6. Functional annotation of the secretome of the saprotrophic basidiomycete *Coprinopsis cinerea* and antimicrobial activity prediction results.**
Secretome functional annotation outputs of emapper (*80*), transmembrane protein prediction by TMBed (*83*) and carbohydrate-active enzyme annotation from dbCAN (*82*), together with the results of antimicrobial activity prediction with AMAPEC.

**Table S7. Functional annotation of the secretome of the saprotrophic ascomycete *Verticillium dahliae* and antimicrobial activity prediction results.**
Secretome functional annotation outputs of emapper (*80*), transmembrane protein prediction by TMBed (*83*) and carbohydrate-active enzyme annotation from dbCAN (*82*), together with the results of antimicrobial activity prediction with AMAPEC.

**Table S8. Description of the 150-fungal genome dataset used for comparative genomic analyses.**
List of the fungal genomes included in the comparative genomic dataset used throughout this study. This dataset is a modified version of a previously studied set of 120 fungal genomes (*85*). This table provides genome identifiers, strain names, assigned lifestyles, metadata about the fungal isolation, genome references, statistics about the genome size and quality, and fungal phylogeny (phylum, class, order).

**Table S9. Annotation and conservation of secreted protein families defined through orthology prediction.**
List of secreted protein families defined through orthology prediction by OrthoFinder (*84*), together with the functional annotation of their most central, representative member (identified with phylorep (*87*)) including transmembrane protein prediction (TMBed (*83*)) and CAZyme annotation (dbcan (*82*)). This table also provides results of antimicrobial activity prediction for protein family representive members which were not annotated as transmembrane or CAZymes.

**Table S10. Antimicrobial activity prediction for secreted proteins of the vascular wilt pathogen *Verticillium dahliae* which were previously studied for their contribution to fungal virulence.**
List of proteins in the secretome of the plant pathogen *Verticillium dahliae* that match (blastp-based identification, sequence identity >95%) proteins registered in PHI-base (*36*) that were previously studied for their contribution to fungal virulence. This table provides results of antimicrobial activity prediction for these proteins, following structure prediction with ESMFold (*74*).

**Table S11. Antimicrobial activity prediction for previously characterized fungal effectors registered in PHI-base.**
List of fungal proteins registered as "effectors (plant avirulence determinant)" in PHI-base (*36*), together with their antimicrobial activity prediction, their closest protein homolog in the 150-genome dataset and their conservation throughout this dataset.

**Table S12. Prediction of antimicrobial activities in LysM effectors annotated in 151 fungal genomes.**
LysM effectors annotated in the set of 150 fungal genomes ((*34*), see Material and Methods for details), as well as in the genome of *Cladosporium fulvum* which secretes the well-studied LysM effector Ecp6 (*37*), listed with their LysM domain identities (as defined by InterPro (*93*)), sequences and results of antimicrobial activity prediction after structure prediction with ESMFold (*74*).

**Table S13. Prediction of antimicrobial activities in the protein family including the AGLIP1 effector protein of the root rot pathogen *Rhizoctonia solani*.**
List of protein homologs of AGLIP1 (effector secreted by *Rhizoctonia solani*) identified and classified in the same family through orthology prediction by OrthoFinder (*84*), together with results of antimicrobial activity prediction following structure prediction with ESMFold (*74*).

**Table S14. Prediction of antimicrobial activities in the protein family including the AVR-Pita effector protein of the rice blast pathogen *Magnaporthe oryzae*.**
List of protein homologs of AVR-Pita (effector secreted by *Magnaporthe oryzae*) identified and classified in the same family through orthology prediction by OrthoFinder (*84*)), together with results of antimicrobial activity prediction following structure prediction with ESMFold (*74*).

**Table S15. Prediction of antimicrobial activities in the protein family including the Vd424Y effector protein of the vascular wilt pathogen *Verticillium dahliae*.**
List of protein homologs of Vd424Y (effector secreted by *Verticillium dahliae*) identified and classified in the same family through orthology prediction by OrthoFinder (*84*), together with results of antimicrobial activity prediction following structure prediction with ESMFold (*74*).

**Table S16. Prediction of antimicrobial activities in the protein family including the VdCP1 effector protein of the vascular wilt pathogen *Verticillium dahliae*.**
List of protein homologs of VdCP1 (effector secreted by *Verticillium dahliae*) identified and classified in the same family through orthology prediction by OrthoFinder (*84*), together with results of antimicrobial activity prediction following structure prediction with ESMFold (*74*).

**Table S17. Compositions of dialysis buffers used for effector protein purification.**
List and composition of dialysis buffers used to purify effector proteins AGLIP1, AVR-Pita, Vd424Y and VdCP1.
